# Global identification of neuronal and astrocytic integral membrane proteins that require Retromer for their endosomal recycling

**DOI:** 10.64898/2026.05.14.724903

**Authors:** Emma Jones, Helena Adams, Kai-en Chen, Fabeeha Maroof, Therese M. Ibbotson, Yasuko Nakamura, Paul J. Banks, Michael D. Healy, Philip A. Lewis, Kate J. Heesom, Brett M. Collins, Kevin A. Wilkinson, Peter J. Cullen, Kirsty J. McMillan

## Abstract

Efficient transport of membrane proteins through the endosomal network is essential for brain development and function, with perturbation implicated in disease. Deficiencies in Retromer, a key regulator of endosomal transport, have been linked to aging-related neurodegenerative disorders including Alzheimer’s and Parkinson’s disease. To better define the neuroprotective role of Retromer, we have applied cell surface restricted proteomics to identify those integral membrane proteins whose recycling to the plasma membrane is mediated by Retromer and associated cargo adaptors, sorting nexin 3 (SNX3), its paralogue sorting nexin 12 (SNX12), and sorting nexin 27 (SNX27) (data available via ProteomeXchange: PXD078277). By comparing primary rat cortical neurons and astrocytes we have identified several cargoes that require either SNX3/SNX12- or SNX27-Retromer complexes for endosomal recycling, including proteins involved in synapse organisation, synaptic signalling and Alzheimer’s disease pathology. We highlight that perturbed Retromer function leads to endosomal enlargement, and we establish a key role of SNX27-Retromer in modulating transport of glutamate across both neuronal and astrocytic membranes via recycling of glutamate transporters EAAT3 (SLC1A1) and EAAT1 (SLC1A3) respectively. Our study provides further mechanistic insight into the consequences of Retromer deficiency for neuronal and astrocytic function, offering new avenues of research in the treatment of neurodegenerative disease.

Graphical Abstract
Suppression of Retromer and the sorting nexins (SNX27, SNX3/SNX12) leads to a significant change in the surface proteome of rat cortical neurons and astrocytes. Focusing on the glutamate transporters, SLC1A1 and SLC1A3, we have validated that SNX27-Retromer is required for their trafficking, with SNX27-Retromer suppression in astrocytes leading to a loss of glutamate uptake.

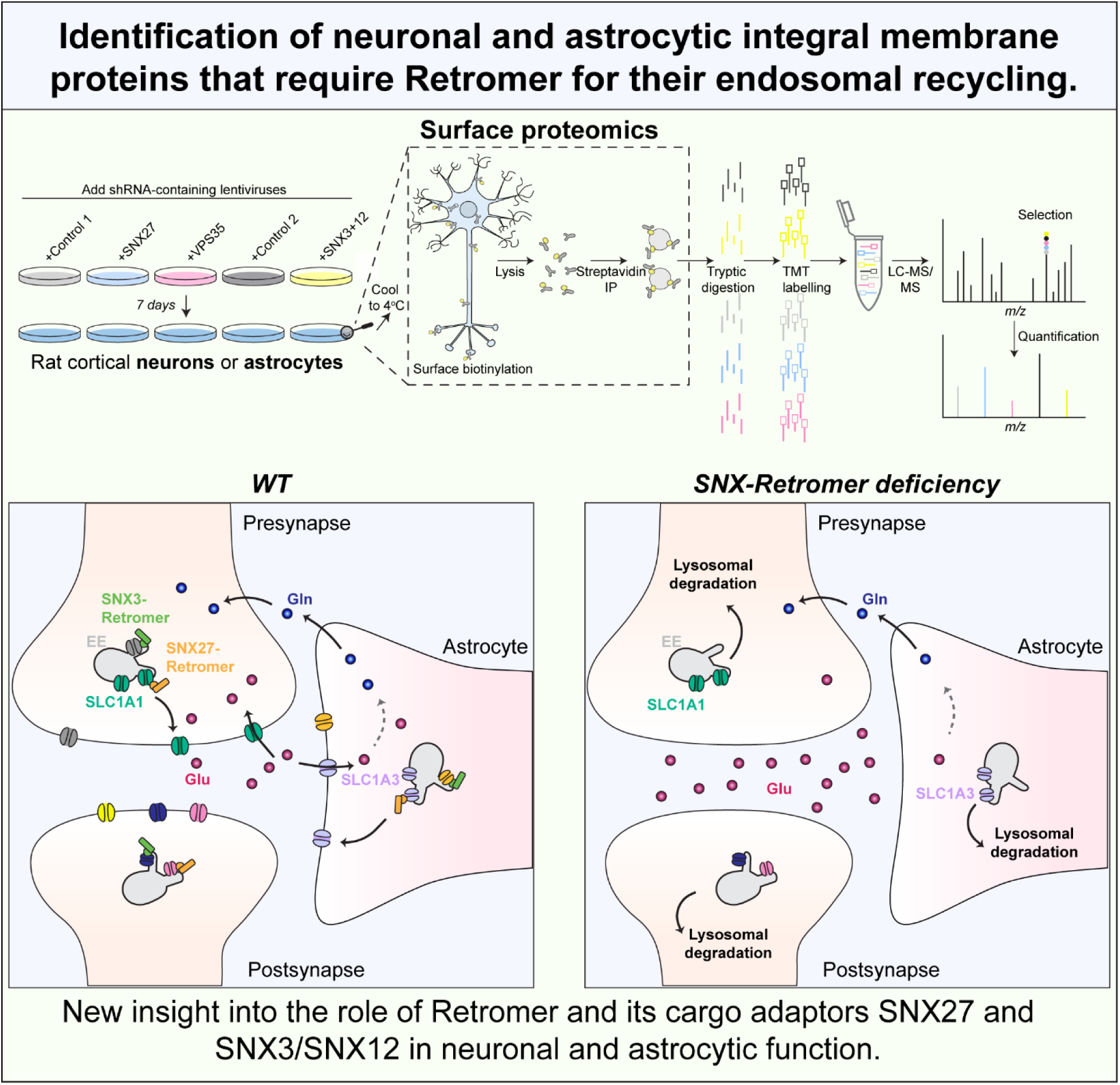

## Introduction

Genetic studies have identified many genes associated with the endosomal network to be causal for numerous neurodegenerative disorders including Alzheimer’s disease, Parkinson’s disease, and Amyotrophic Lateral Sclerosis [1–7]. These studies, alongside clinical evidence [8–14], highlight the endosomal network as a key biological pathway affected in neurodegenerative pathology. The endosomal network comprises a series of intracellular membrane-bound compartments that controls the transport of integral membrane proteins, and their associated proteins and lipids (termed cargoes), to a variety of cellular compartments, a role that is vital for cell function and survival. On entering the network, cargoes essentially have two fates: they are either sorted towards the lysosome for degradation, or are retrieved and recycled to the plasma membrane, the Golgi apparatus or other specialized organelles [15–19].

Human Retromer is a stable heterotrimer of vacuolar protein sorting protein 35 (VPS35), either VPS26A or VPS26B [20, 21], and one of three VPS29 isoforms [22, 23], and regulates the endosomal retrieval and trafficking of numerous cargoes away from the degradative pathway [23]. To function, Retromer associates with accessory proteins and cargo adaptors including sorting nexins (SNXs) [24–34], the WASH complex [35–39], RAB GTPase regulators including TBC1D5 and ANKRD27/VARP [40–46], and RME8 [47]. Retromer expression is significantly decreased in neurodegenerative patients [10–14, 48], with stabilisation of Retromer expression being neuroprotective [49–52]. Disease causing mutations in Retromer have been identified in patients, including the VPS35(D620N) mutation which causes an autosomal dominant form of Parkinson’s disease [53, 54] and the *de novo* mutation VPS35(L625P) which has been associated with early onset Alzheimer’s disease [55]. In addition, neuronal endosomal swelling has been classed as an early hallmark of Alzheimer’s disease indicative of a perturbation in endosomal cargo sorting [8, 9, 56]. This has led to Retromer, and more generally the endosomal network, being identified as potential therapeutic targets in the treatment of neurodegenerative disorders.

Sorting nexin-27 (SNX27) is an important Retromer cargo adaptor that contains an amino-terminal PDZ (PSD-95, Disc-large and ZO-1) domain, a PX (Phox) domain that enables direct binding to phosphatidylinositol 3-monophosphate (PtdIns(3)P3) present on the endosomal membrane, and a carboxy-terminal FERM (4.1/ezrin/radixin/moesin) domain. Through its PDZ domain, SNX27 can simultaneously interact with Retromer and cargoes containing a PDZ binding motif (PDZbm) in their carboxy-terminal tail [26, 29, 31, 57]. Through this dual interaction SNX27 regulates the retrieval and recycling of hundreds of cargoes to the plasma membrane [26, 29, 31, 58, 59]. Through its FERM domain, SNX27 may also associate with cargoes containing a NPxY or NxxY sorting motif (x refers to any residue), with SNX27 showing a preference for sequences that are phosphorylated at the tyrosine residue [29, 60, 61]. SNX27 also interacts with the SNX-BAR (Bin/Amphiphysin/Rvs) proteins SNX1 and SNX2 [27, 28, 31, 62] and the WASH complex [37] to couple sequence-dependent retrieval of SNX27 cargoes with tubule formation during the process of cargo recycling. Importantly, SNX27 dysfunction is implicated in neurodegeneration and Down’s syndrome, which greatly increases the risk for developing Alzheimer’s disease [63–65].

An additional cargo adaptor complex essential for the sequence-dependent retrieval and recycling of cargoes is the sorting nexin-3 (SNX3)-Retromer assembly [25, 30, 66, 67]. SNX3 also has a PX domain and, alongside Rab7A, has been shown to be important for facilitating Retromer’s localisation to the endosome [1, 25, 68]. SNX3 directly binds to a subset of cargo containing the poorly defined sequence motif [+/-]-x-∅-x-[L/M] (∅ - bulky aromatic residue) with many of these cargoes trafficked through the retrograde endosome-to-TGN pathway [25, 30, 66, 67]. Moreover, SNX3-Retromer has also been implicated in regulating the expression of proteins present at the cell surface [30, 69–71] and single nucleotide polymorphisms (SNPs) in SNX3 are associated with Alzheimer’s disease [1]. Of note, humans express a SNX3 homologue, SNX12, which shares a 79.5% protein sequence identity [72]. SNX12 has also been implicated in Alzheimer’s disease through modulation of amyloid precursor protein (APP) processing via an interaction with BACE1 [73]. Moreover, SNX3 and SNX12 depletion have been shown to affect neurite outgrowth [74–76].

A more complete definition of the range of neuronal cargoes dependent on the endosomal system is vital to further our understanding of how endosomal dysfunction leads to neurodegenerative pathology. Here, we have utilised quantitative proteomic approaches in primary rat cortical cultures to identify integral membrane proteins that depend on the SNX27-Retromer and SNX3/SNX12-Retromer complexes for their surface expression in neurons. Given the evolving appreciation of the role of neuronal supporting cells in the initiation and progression of neurodegenerative disease we have also extended our methodological approach through a parallel analysis of these sorting complexes in astrocytes. Our global analysis establishes the role of these complexes in neuronal and astrocytic function, particularly in glutamate uptake, and highlights many proteins and pathways whose perturbation may drive astrocytic and neuronal dysfunction, neuronal death and ultimately neurodegenerative disease.

## Results

### Retromer, SNX27 and SNX3/SNX12 suppression significantly alters the surface proteome of primary rat cortical neurons

To identify neuronal cargoes that depend on SNX27-Retromer or SNX3-Retromer for their surface expression we transduced DIV12 rat cortical neuronal cultures with lentiviruses co-expressing GFP and shRNAs targeting *Vps35* (core component of Retromer), *Snx27* or *Snx3*, or a control non-targeting shRNA (Control 1) [59, 77]. To prevent redundancy between SNX3 and SNX12 due to their high level of sequence identity, we also generated a lentivirus co-expressing mCherry and either a shRNA targeting *Snx12* or the control non-targeting shRNA. The *Snx12* shRNA was transduced alongside the *Snx3* shRNA and compared to an equivalent control condition using two non-targeting shRNAs co-expressed with either GFP or mCherry (Control 2). After 7 days we observed a significant suppression of VPS35 (> 80%), SNX27 (> 80%), SNX3 (> 80%) and SNX12 (> 65%) across the three biological repeats **(Figures 1A-B)**. By suppressing protein expression rather than using a knockout (KO) model our approach is physiologically relevant to neurodegenerative patients where Retromer levels are reduced, in contrast with Retromer KO which is embryonically lethal [78]. As previously reported, VPS35 suppression led to endosomal swelling in the rat cortical neurons [79], with enlarged endosomes also found in our SNX27 and SNX3/SNX12 suppressed neurons, indicative of a trafficking defect (**Supplementary Figure 1A**). As SNX3 has previously been shown to contribute to Retromer recruitment onto endosomal membranes [1, 25], we also measured VPS35 co-localisation with SNX6 as a marker of the endosomal recycling subdomain. SNX3/12 suppression led to a significant decrease in co-localisation of VPS35 with SNX6 (**Supplementary Figure 1B-C**).

**Figure 1.**
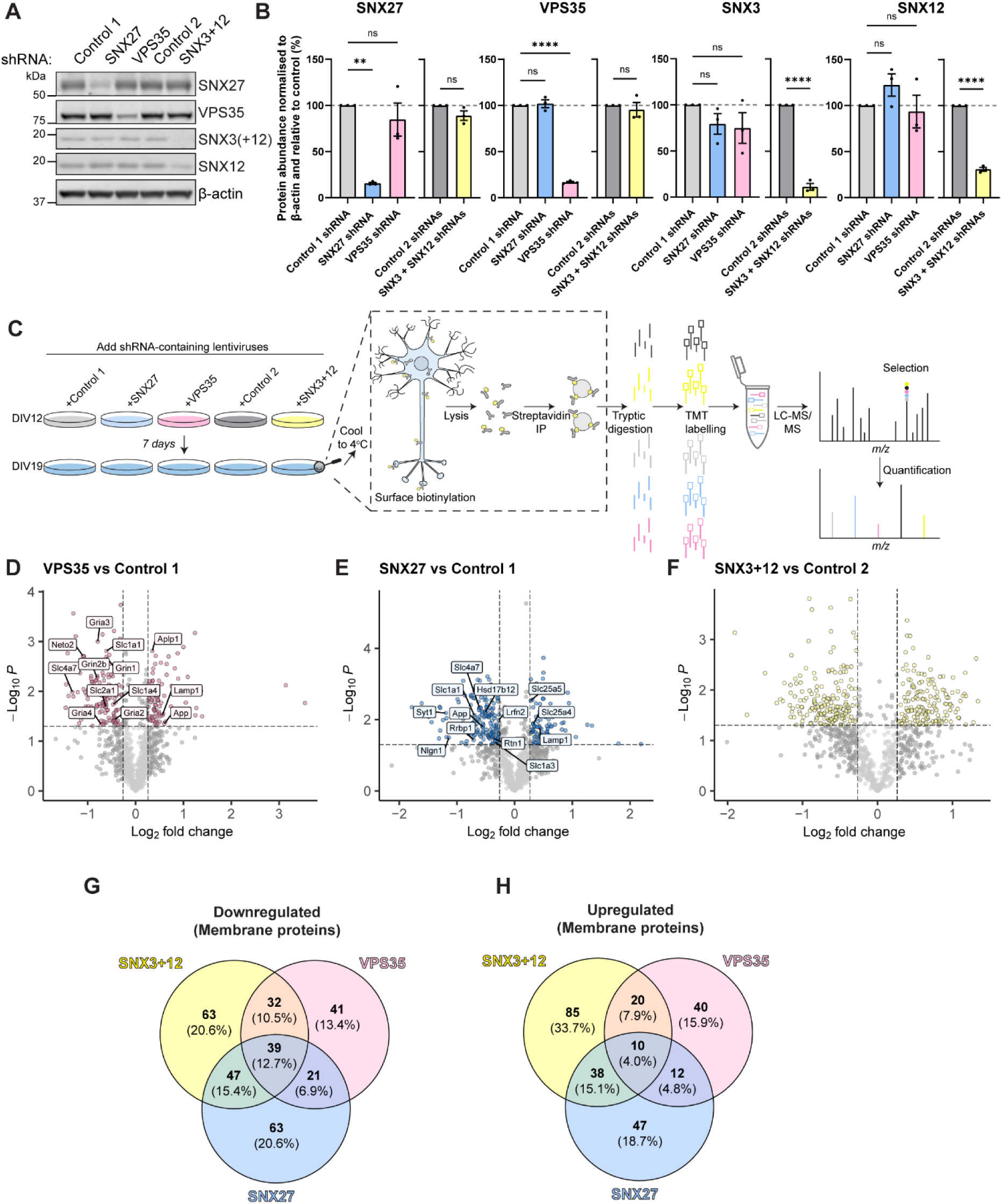
Retromer, SNX27 and SNX3+SNX12 suppression significantly alters the surface proteome of primary rat cortical neurons. **(A)** Fluorescence-based western blot of DIV19 rat cortical neurons transduced with either control non-targeting shRNAs (Control 1 and 2) or shRNAs against SNX27, VPS35 or SNX3+SNX12. Blotted for endogenous VPS35, SNX27, SNX3 and SNX12 with β-actin used as a protein load control. **(B)** Quantification from western analyses of *(A)* from three independent experiments (n = 3). Data expressed as a percentage of the relevant control shRNA. SNX27 and VPS35 shRNA data analysed by an one-way ANOVA followed by a post hoc Dunnett’s test compared to Control 1. SNX3+SNX12 shRNA data analysed by an unpaired t-test compared to Control 2. Error bars represent mean ± SEM. ****, p ≤ 0.0001; **, p ≤ 0.01; ns, not significant. **(C)** Schematic depicting the biotinylation and isolation of surface membrane proteins in DIV19 rat cortical neuronal cultures followed by TMT-based quantitative proteomics. **(D-F)** TMT surfaceome of **(D)** VPS35 suppressed, **(E)** SNX27 suppressed or **(F)** SNX3+SNX12 suppressed rat cortical neurons compared to non-targeting control shRNAs (Control 1 or Control 2). Data quantified across three independent experiments (n = 3). Plotted proteins are membrane proteins present in all three data sets and analysed using a one-sample t-test. Vertical grey line represents the threshold for depleted or enriched proteins (log2 fold change +/- 0.26) compared to the relevant control. Horizontal grey line represents the threshold for statistical analysis (p ≤ 0.05). Coloured circles are the proteins that are significantly changed compared to the relevant non-targeting shRNA control. Highlighted proteins are previously validated cargoes. **(G-H)** Venn diagrams comparing the number of membrane proteins significantly **(G)** downregulated or **(H)** upregulated at the surface of rat cortical neuronal cultures after treatment with shRNAs against VPS35, SNX27 and SNX3+SNX12.

Using the shRNA treated rat cortical cultures, we isolated the surface proteins by performing surface restricted biotinylation followed by streptavidin affinity isolation. The efficiency of the surface biotinylation was validated by western blotting, showing an enrichment of the surface protein N-cadherin, which was unaffected after VPS35, SNX27 or SNX3/SNX12 suppression, and a lack of labelling of the abundant cytosolic protein β-actin (**Supplementary Figure 2A**). Isolated proteins were digested followed by tandem mass tagging (TMT) and resulting peptides were quantitatively identified using liquid chromatography-tandem mass spectrometry across three independent biological repeats **(Figure 1C)**. Data were normalised to total peptide amount to provide relative changes to the surface proteome.

For the VPS35 suppressed neuronal cultures of a total of 5214 quantified proteins, 133 integral proteins were significantly depleted (log2 fold change −0.26, p < 0.05) and 82 membrane proteins were enriched (log2 fold change +0.26, p < 0.05) when compared with the control 1 shRNA treated neuronal cultures **(Figure 1D, Supplementary Table 1)**. Validating our approach, downregulated proteins included the well-established Retromer cargo glucose transporter-1 (GLUT1/SLC2A1) [29], the ionotropic glutamate AMPA receptors (GRIA2, GRIA3, GRIA4) and NMDA receptors (GRIN1, GRIN2b) [80–85], as well as many other previously identified proteins including the sodium bicarbonate cotransporter-3 (SLC4A7) (**Figure 1D, Supplementary Table 2**) [29, 79]. SLC4A7 surface expression was also decreased by SNX27 and SNX3/SNX12 suppression which we validated biochemically using surface biotinylation, streptavidin affinity capture coupled with quantitative western analysis (47% reduction after SNX27 suppression (p = 0.0001), 82% reduction after VPS35 suppression (p < 0.0001), n = 3, one-way ANOVA) and a 70% reduction after SNX3/SNX12 suppression (p = 0.0012, n = 3, unpaired t-test) (**Supplementary Figure 2B-C**). We also found a significant decrease in total SLC4A7 expression after SNX27 suppression (63% reduction; p = 0.0091; n = 3, one-way ANOVA) and VPS35 suppression (56% reduction; p = 0.0152; n = 3, one-way ANOVA). SNX3/SNX12 suppression also led to a 36% reduction in total SLC4A7 levels (p = 0.0264; n = 3, unpaired t-test) suggesting that mis-trafficked SLC4A7 undergoes degradation in the lysosome.

For SNX27 suppressed neuronal cultures, 170 integral proteins were depleted (log2 fold change −0.26, p ≤ 0.05) and 107 integral proteins were enriched (log2 fold change +0.26, p ≤ 0.05) compared with the control 1 shRNA treated neuronal cultures **(Figure 1E, Supplementary Table 1).** In previous work, we performed a SNX27 interactome analysis in rat neuronal cortical cultures [59]. Comparing these two data sets, we found 9 membrane proteins that were significantly affected by SNX27 suppression and were also present in the SNX27 interactome making them high-confidence cargoes (**Figure 1E, Supplementary Table 3**). Of the depleted proteins these included the synaptic adhesion protein LRFN2, the glutamate transporter SLC1A3 (EAAT1/GLAST), and the sodium bicarbonate cotransporter-3 SLC4A7, all of which contain a Type I PDZbm in their carboxy terminal tail. In previous work we have validated that LRFN2 and SLC1A3 can interact with SNX27 directly through their PDZbm [59]. Moreover, of the 170 depleted proteins in the SNX27 suppressed neuronal cultures 46 proteins contained a PDZbm indicative of binding to SNX27. We ran the last 10 amino acids of these proteins against the SNX27-PDZ in AlphaFold2 and took the average interface predicted template modelling score (iPTM) from 5 models. 42 of the proteins had an iPTM value higher than 0.6 which predicts high-confidence binding (**Supplementary Table 4**).

For the SNX3/SNX12 suppression, 181 integral proteins were depleted (log2 fold change −0.26, p ≤ 0.05) and 153 integral proteins were enriched (log2 fold change +0.26, p ≤ 0.05) compared with the control 2 shRNA treated neuronal cultures **(Figure 1F, Supplementary Table 1).** To identify which integral proteins depleted from the surface are likely to be a direct consequence of binding to SNX3-Retromer, we screened the 25 most depleted single-pass transmembrane proteins with annotated cytoplasmic tail sequences for any predicted association with SNX3, VPS26A and VPS35 using AlphaFold2 and AlphaFold3 (**Supplementary Table 5**). Likely due to the flexibility in the binding motif of SNX3-Retromer cargo, there was discrepancy between confidence of predictions across models assessed, however, this analysis highlighted a high confidence association of ADAM10 with the same interface of SNX3 and VPS26A previously structurally characterised in complex with DMT1-II [66] (**Supplementary Table 5**). The motif of association is consistent with the previously published consensus motif for SNX3 (E-s-Y-q-M; [+/-]-x-∅-x-[L/M]) (**Supplementary Figure 3A**). Although further investigation is required to validate these predicted interfaces, this provides evidence that select proteins depleted from the surface with SNX3/12 suppression are likely to be the direct result of reduced SNX3/12-Retromer mediated endosomal recycling. However, as SNX3/12 suppression led to a significant decrease in co-localisation of VPS35 with SNX6 (**Supplementary Figure 1B-C**), any proteins depleted from the surface in this condition could also be a result of reduced Retromer residency on endosomes.

Comparing across the different suppression conditions, 60 integral membrane proteins were decreased by both SNX27 and VPS35 suppression whilst 71 proteins were decreased by both SNX3/SNX12 and VPS35 suppression. Of these, 39 proteins were shared across all conditions (**Figure 1G and Supplementary Figure 3B)**. For the enriched proteins there were 22 integral proteins enriched by both SNX27 and VPS35 suppression and 48 proteins enriched by both SNX3/SNX12 and VPS35 suppression. Of these only 10 proteins were shared across all conditions (**Figure 1H and Supplementary Figure 3C**).

### Retromer suppression affects multiple key neuronal pathways

To analyse our proteomic data further we focused initially on the surface proteome of the VPS35 suppressed neuronal cultures. Importantly, of the 215 proteins that were significantly altered on the surface following VPS35 suppression, 136 of these have not been previously described (**Supplementary Table 2**). Gene ontology using Cluster Profiler highlighted a range of proteins involved in important neuronal pathways that were significantly affected following suppression of VPS35. The most significant downregulated pathways included proteins involved in regulating presynaptic membrane potential, synaptic adhesion, synaptic transmission, synapse structure and organisation **(Figure 2A)**. Enriched pathways included proteins involved in synapse assembly, exocytosis, APP processing and amyloid-β response (**Figure 2B**). We examined the proteins within these VPS35 GO pathways and analysed their expression after VPS35, SNX27 or SNX3/SNX12 suppression. In addition to AMPA and NMDA receptors, our screen highlighted the glutamate pathway as a key pathway affected by Retromer suppression. Kainate receptors, metabotropic glutamate receptors and glutamate transporters (including SLC1A1/EAAT3 and SLC1A2/EAAT2/GLT1) were all found to be significantly reduced at the neuronal surface after VPS35 suppression (**Figure 2C**). Glutamate is the main excitatory neurotransmitter, with perturbation of this system leading to several neurological disorders. Moreover, many of the proteins involved in the GABAergic pathway, including several subunits of the ionotropic GABA type A receptors (GABA_A_Rs), were also significantly reduced at the surface of VPS35 suppressed neurons (**Figure 2C**). This is of interest as disturbances in the glutamatergic and GABAergic systems and an imbalance in the excitatory-inhibitory pathways are heavily implicated in neurodegenerative pathology [86–88]. SNX27 and SNX3/SNX12 suppression also caused a significant loss of proteins involved in the glutamatergic and GABAergic pathways (**Figure 2C**). Surprisingly, AMPA receptors were not reproducibly lost from the neuronal surface of our SNX27 suppressed cultures. We previously showed that AMPA receptors are indirectly affected by SNX27 through the trafficking of adhesion molecules including LRFN2 [59]. Importantly, our surface proteomics showed a significant loss of LRFN2 after SNX27 suppression which independently corroborates our previous findings (**Figure 2C**). The loss of LRFN2 however was only by 18% suggesting that it may be necessary to suppress SNX27 expression even further to see an effect on AMPA receptor expression [59]. Surprisingly, VPS35 suppression resulted in an increase in LRFNs (LRFN1, 2 and 4) at the neuronal surface (**Figure 2C**) suggesting that SNX27 can act independently to Retromer in the trafficking of a sub-set of cargoes, potentially through the ESCPE-1 complex [27, 28]. Indeed, we found a decrease of 63 membrane proteins after SNX27 suppression independently of Retromer or SNX3/SNX12 (**Figure 1G and Supplementary Table 1**).

**Figure 2.**
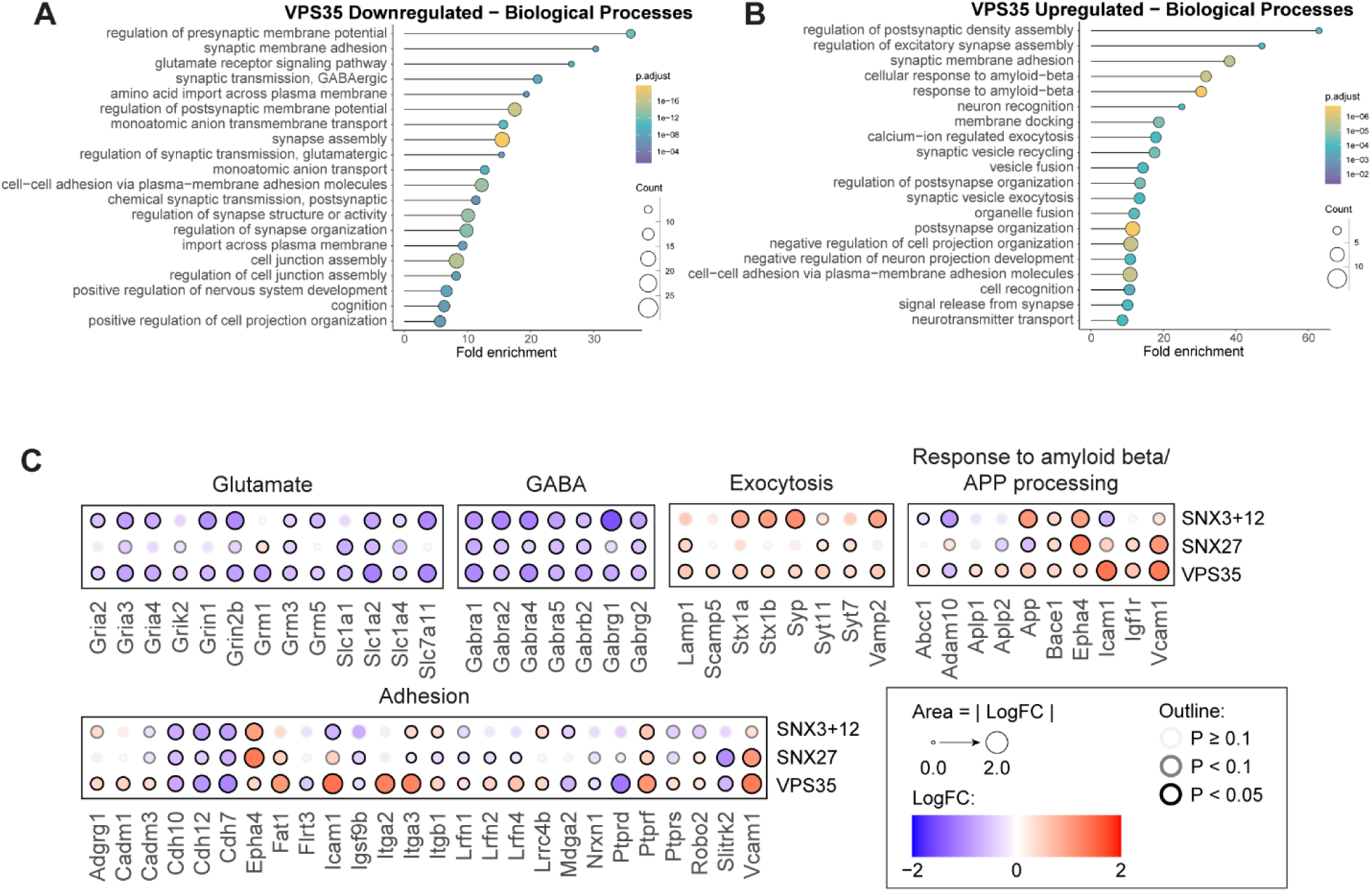
Retromer suppression affects multiple key neuronal pathways. **(A)** Downregulated or **(B)** upregulated surface proteins in VPS35 suppressed rat cortical neurons converge into biological pathways as shown by gene ontology analyses using Cluster Profiler. **(C)** Dot-plots representing log_2_ fold change and p-value in quantified abundances of proteins separated into key biological pathways after VPS35, SNX27 or SNX3+SNX12 suppression. This includes the glutamatergic and GABAergic pathways, adhesion, exocytosis, APP processing and amyloid-beta response.

In previous work we reported an increase in APP surface expression in VPS35 KO H4 cells and hypothesised that this was due to an increase in lysosomal exocytosis [79]. Mutations in APP are causal for Alzheimer’s disease with abnormal processing of APP and the generation of toxic amyloid-β aggregates hallmarks of Alzheimer’s pathology [89]. We corroborated these data in neurons showing an increase in the surface expression of APP after VPS35 suppression (**Figure 1D**, **Figure 2C**). We also validated this biochemically using surface biotinylation, streptavidin affinity capture coupled with quantitative western analysis. Both VPS35 suppression and SNX3/SNX12 suppression showed a significant increase of APP at the cell surface (52% enrichment after both VPS35 suppression (p < 0.0001, n = 4, one-way ANOVA) and SNX3/SNX12 suppression (p = 0.0173, n = 4, unpaired t-test) (**Supplementary Figures 2B-C)**. APP is cleaved by α- or β-secretases to produce a low molecular weight C-terminal fragment (CTF). We found a significant increase in the cell surface expression of the APP CTF (45% enrichment after VPS35 suppression (p = 0.0002, n = 4, one-way ANOVA); 102% enrichment after SNX3/SNX12 suppression (p = 0.0011, n = 4, unpaired t-test). We also found a significant increase in the surface expression of the lysosomal-associated membrane protein-1 (LAMP-1) after VPS35 suppression in our proteomic data indicating an increase in lysosomal exocytosis (**Figure 1D**, **Figure 2C**). Moreover, we found an increase in surface expression of vesicle-associated membrane protein 2 (VAMP/synatobrevin-2), syntaxin-1A (STX1A) and synaptotagmin-7 (SYT7) (**Figure 2C**), all of which have been associated with lysosomal exocytosis [90, 91], further highlighting an important role for Retromer in maintaining lysosomal health. Whilst we did see a trend towards an increase in surface LAMP1 expression by western analysis after VPS35 and SNX27 suppression, the data did not reach significance. However, SNX3/SNX12 suppression caused a significant increase of LAMP1 at the surface by 102% (p = 0.0210, n = 4, unpaired t-test) **(Supplementary Figures 2B-2C)**.

In addition to APP, we found an increase in the surface expression of several proteins involved in APP processing and amyloid-β response in the VPS35 suppressed neuronal cultures. These included β-secretase 1 (BACE1), [92], APP-like β-secretase substrates APLP1 and APLP2 [93], transporter ABCC1 [94–96], ephrin type-A receptor 4 (EPHA4) [97, 98], and insulin-like growth factor-1 (IGFR1) [99], and Synaptophysin (SYP) [100] (**Figure 2C**). Interestingly, ADAM10, an α-secretase involved in APP cleavage, was also significantly lost at the surface of VPS35 suppressed neuronal cultures. ADAM10 cleaves APP in a non-amyloidogenic manner [101, 102]. It is therefore interesting to speculate whether a reduction in ADAM10 at the surface of neurons would lead to an increase in amyloid-β peptide production. Whilst Retromer has been shown to be important in APP trafficking [55, 103–105], our data highlights several other proteins and pathways through which Retromer can also influence APP processing and amyloid-β processing and response.

Taken together, our proteomic analysis highlights several essential neuronal pathways affected by Retromer, SNX27 and SNX3/SNX12 dysfunction including the glutamatergic and GABAergic pathways, APP processing, amyloid-β uptake and lysosomal exocytosis. Furthermore, GO analysis highlights many key adhesion proteins involved in synaptic assembly and organisation (**Figure 2C**). Future work will be essential to address how perturbations of these proteins and pathways can lead to dysregulated synaptic health and its implication for neuronal death and neurodegenerative disease.

### Role of Retromer, SNX27 and SNX3/SNX12 in Astrocytes

Previous work investigating the role of the endosomal network and Retromer in neurodegeneration has focused primarily on neurons. Astrocytes are known to play a vital role in neuronal health and neurodegeneration but the role of the endosomal network and Retromer in astrocyte function is unclear and understudied, leaving a critical gap in our knowledge. To assess the astrocytic proteins that depend on Retromer for their surface expression we produced astrocyte enriched cultures with over 80% of cells GFAP positive **(Figure 3A-B)**. These cells also expressed the glutamate transporter SLC1A3/GLAST, and the calcium binding protein S100β (**Figure 3A**), both of which are primarily expressed in astrocytes. Using our shRNAs we transduced these astrocytic cultures for 7 days acquiring a sufficient level of suppression of VPS35 (>90%), SNX27 (>90%), SNX3 (>90%) and SNX12 (>70%) across the three biological repeats **(Supplementary Figure 4A-B)**. Whilst these shRNAs are specific to their targets, in astrocytes, we did find a small but significant decrease of SNX27 expression by 15% after SNX3/SNX12 suppression.

**Figure 3.**
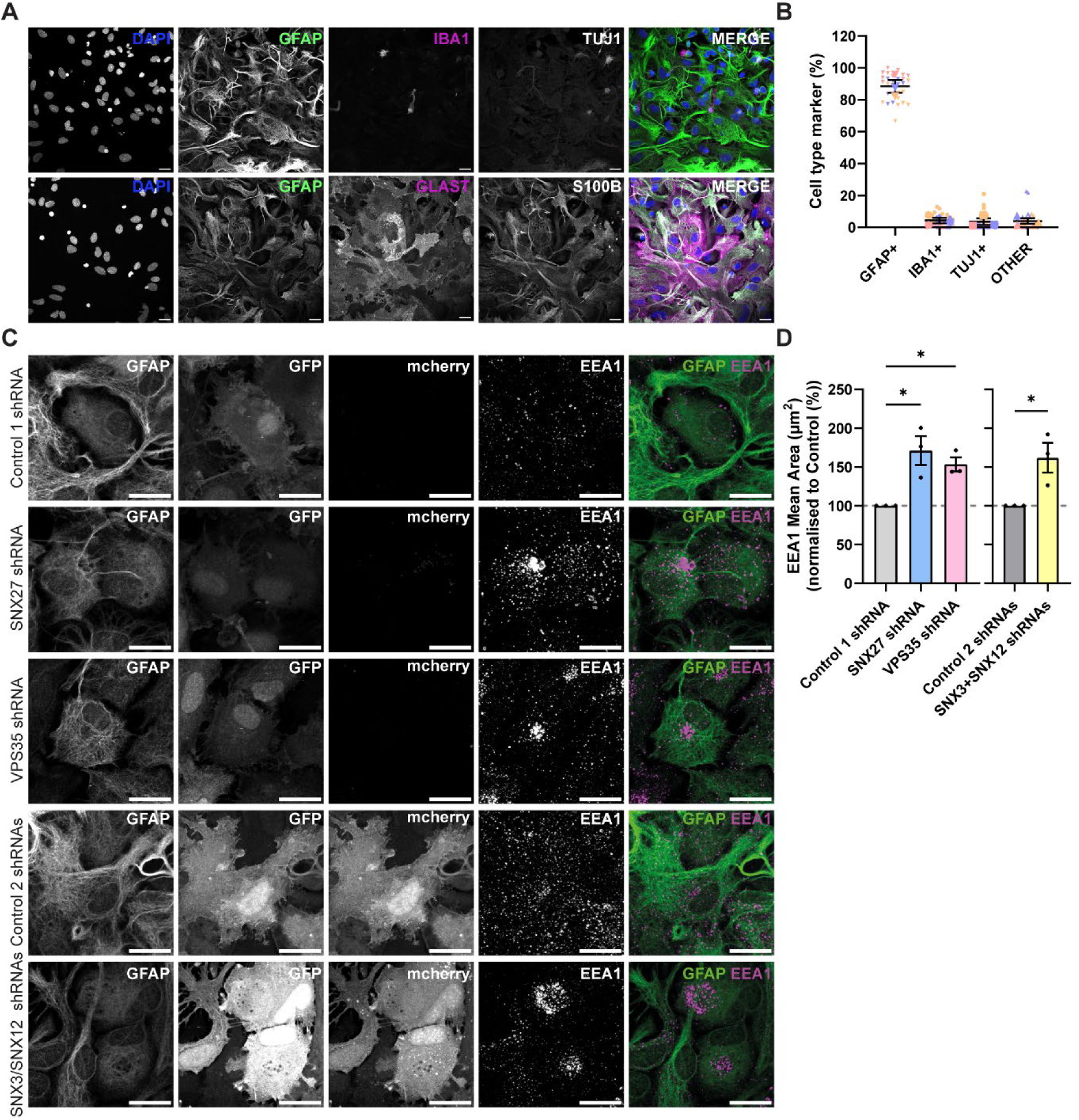
Retromer, SNX27 or SNX3+SNX12 suppression in astrocytes leads to endosomal swelling. **(A)** Immunofluorescence images of astrocyte markers GFAP (green), SLC1A3 (pseudo coloured in magenta) and S100β (pseudo coloured in white) (bottom panel), the microglial marker IbaI (pseudo coloured in magenta) and the neuronal marker TUJ1 (pseudo coloured in white) (top panel) and astrocyte markers GFAP (green), in rat cortical astrocyte cultures. Scale bars, 20 µm. **(B)** Quantification of number of GFAP positive astrocyte cells, IbaI positive microglial cells and TUJ1 positive neuronal cells in rat cortical astrocyte cultures from three independent experiments (n = 3, quantified across 30 images and 1,585 cells). Data expressed as a percentage of the total number of cells counted using DAPI. **(C)** Immunofluorescence images of early endosome marker EEA1 (pseudo coloured in magenta) in astrocytes labelled with GFAP (pseudo coloured in green) in shRNA transduced astrocyte cultures (shRNAs with GFP/mCherry). Scale bars, 20 µm. **(D)** Quantification of the EEA1 mean area (μm^2^) in astrocytes transduced with either control non-targeting shRNAs (Control 1 and 2) or shRNAs against SNX27, VPS35 or SNX3+SNX12 from three independent experiments (n = 3, quantified across 30 images, > 118 cells quantified per condition). Data expressed as a percentage of the relevant control shRNA. SNX27 and VPS35 shRNA data analysed by an one-way ANOVA followed by a post hoc Dunnett’s test compared to Control 1. SNX3+SNX12 shRNA data analysed by an unpaired t-test compared to Control 2. Error bars represent mean ± SEM. *, p ≤ 0.05.

Interestingly, similar to the neurons we found a significant increase in the size of the endosomes in our VPS35 (53% increase, p = 0.0349, n = 3, one-way ANOVA), SNX27 (71% increase, p = 0.0102, n = 3, one-way ANOVA) and SNX3/SNX12 (62% increase, p = 0.0317, n = 3, unpaired t-test) suppressed astrocytes compared to the relevant non-targeting shRNA treated controls **(Figure 3C-D**). We again performed surface proteomics using surface biotinylation experiments followed by streptavidin affinity isolation, TMT labelling and liquid chromatography-tandem mass spectrometry across three independent biological repeats. Of a total of 4609 quantified proteins, 137 membrane proteins were depleted (log2 fold change −0.26, p ≤ 0.05) and 56 membrane proteins were increased at the surface after VPS35 suppression (log2 fold change +0.26, p ≤ 0.05) (**Supplementary Figure 4C, Supplementary Table 6),** whilst 80 membrane proteins were depleted (log2 fold change −0.26, p ≤ 0.05) and 114 membrane proteins were increased (log2 fold change +0.26, p ≤ 0.05) after SNX27 suppression **(Supplementary Figure 4D, Supplementary Table 6)** compared with the control 1 shRNA treated cells. 22 proteins depleted on the surface with SNX27 suppression were predicted by AlphaFold2 to interact with SNX27 PDZ domain via their PDZ binding motif (**Supplementary Table 7**). For the SNX3/SNX12 suppressed astrocyte cultures 116 membrane proteins were depleted (log2 fold change −0.26, p ≤ 0.05) and 27 membrane proteins were increased (log2 fold change +0.26, p ≤ 0.05) compared with the control 2 shRNA treated astrocyte cultures **(Supplementary Figure 4E, Supplementary Table 6).** Analysis of the cytosolic tails of surface depleted proteins with SNX3/12 suppression using AlphaFold2 and AlphaFold3 also predicted binding of additional cargo proteins alongside ADAM10, including PTGFRN and NRP2 **(Supplementary Table 8)**. Of note, this analysis also predicted the binding interface of SNX3-Retromer with the transferrin receptor (TFRC), previously shown to be a cargo of this pathway [70]. Comparing across the different suppression conditions, 60 membrane proteins were decreased by both SNX27 and VPS35 suppression whilst 61 proteins were decreased by both SNX3/SNX12 and VPS35 suppression suggesting that there are a similar number of cargoes for each Retromer-SNX complex. Of these, 31 proteins were shared across all conditions (**Figure 4A and Supplementary Figure 4F)**. For the increased proteins there were 30 proteins enriched at the surface after SNX27 and VPS35 suppression and 8 proteins enriched at the surface after SNX3/SNX12 and VPS35 suppression. Of these only 5 proteins were shared across all conditions (**Figure 4B and Supplementary Figure 4G**).

**Figure 4.**
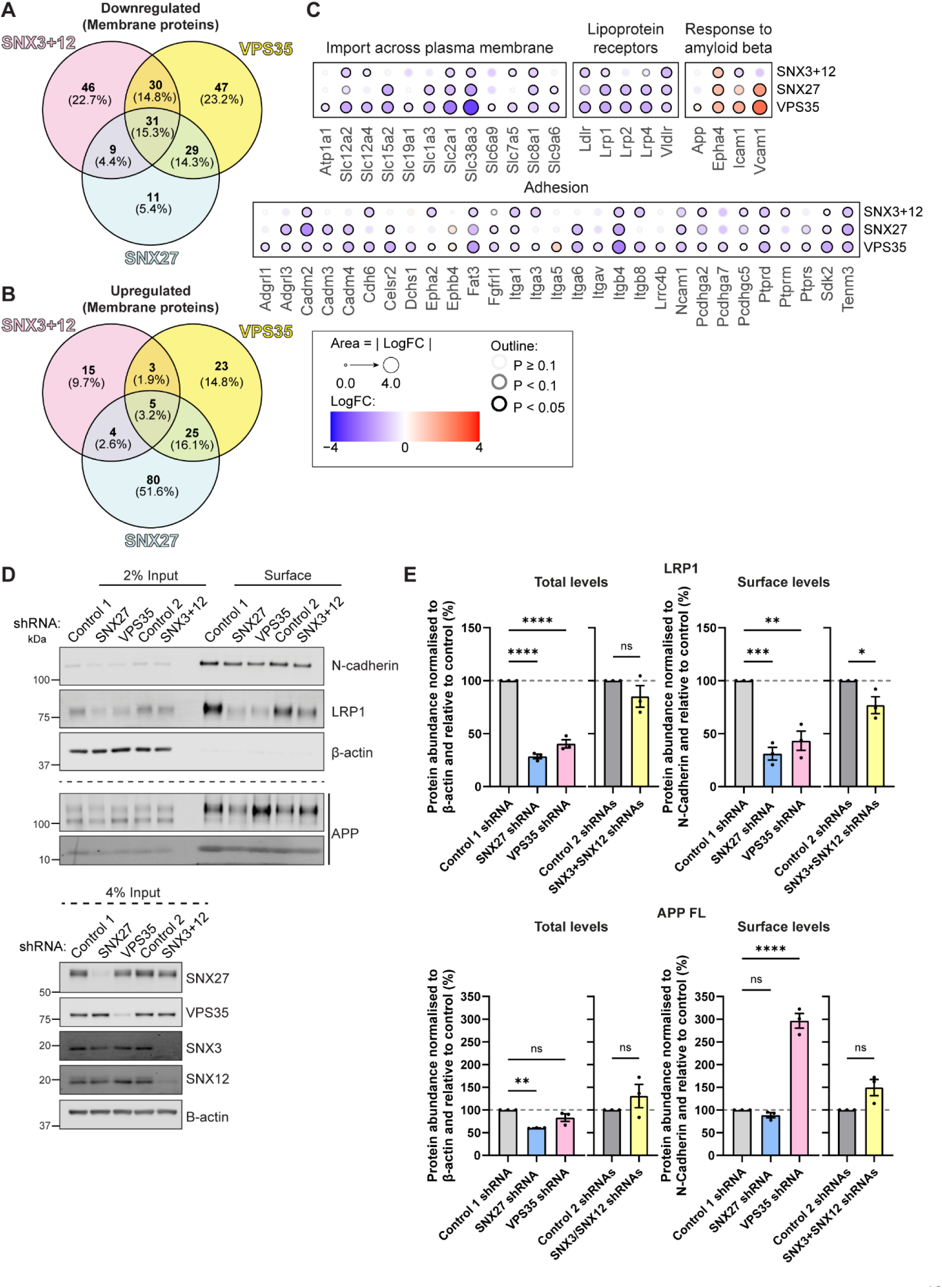
Retromer, SNX27 or SNX3+SNX12 suppression significantly alters the surface proteome of primary rat cortical astrocytes. **(A-B)** Venn diagrams comparing the number of membrane proteins significantly **(A)** downregulated or **(B)** upregulated at the surface of rat cortical astrocyte cultures after treatment with shRNAs against VPS35, SNX27 or SNX3+SNX12. **(C)** Dot-plots representing log_2_ fold change and p-value in quantified abundances of proteins separated into key biological pathways after VPS35, SNX27 or SNX3+SNX12 suppression. This includes the lipoprotein receptors and proteins involved in amyloid-beta response, adhesion and import across the membrane. **(D)** Fluorescence-based western analysis after surface biotinylation and streptavidin agarose capture of membrane proteins of rat cortical astrocytes transduced with control non-targeting shRNAs (Control 1 and 2) or shRNAs against SNX27, VPS35 or SNX3+SNX12. Endogenous surface and total levels of N-Cadherin, LRP1 and APP are shown. The expression level of VPS35, SNX27 and SNX3/SNX12 are also shown with β-actin used as a protein load control. **(E)** Quantification from western analyses of *(D)* from three independent experiments (n = 3). Surface LRP1 and APP are normalised to control N-Cadherin whilst total levels are normalised to β-actin. Data expressed as a percentage of the relevant control shRNA. SNX27 and VPS35 shRNA data analysed by an one-way ANOVA followed by a post hoc Dunnett’s test compared to Control 1. SNX3+SNX12 shRNA data analysed by an unpaired t-test compared to Control 2. Error bars represent mean ± SEM. ****, p ≤ 0.0001; ***, p ≤ 0.001; **, p ≤ 0.01; *, p ≤ 0.005; ns, not significant.

GO analysis revealed that the most significantly downregulated pathways after VPS35 suppression in astrocytes included proteins involved in adhesion, transport across the membrane, and synapse assembly and organisation **(Supplementary Figure 4H)**. Similar to the neuronal cultures, GO highlighted proteins involved in APP processing and amyloid-β response to be significantly upregulated in the VPS35 suppressed astrocyte cultures, including an increase in surface APP expression (**Figure 4C, Supplementary Figure 4I**). Surface biotinylation, streptavidin affinity capture coupled with quantitative western analysis validated that VPS35 suppression caused a significant increase in APP surface expression by 197% (p < 0.0001, n = 3, one-way ANOVA). There was also evidence of an increase in the surface expression of the APP CTF but in astrocytes the protein level was difficult to quantify (**Figure 4D)**. SNX3/SNX12 also showed a trend towards an increase in APP surface levels (p = 0.0502, n = 3, unpaired t-test). Other upregulated pathways after VPS35 suppression highlighted by GO analysis included migration and proliferation **(Supplementary Figure 4I)**.

Analysing the proteins within the GO pathways further, we found a significant depletion of several members of the low-density lipoprotein receptor family including LDLR, VLDLR, LRP1, LRP2 and LRP4 in the VPS35 suppressed astrocyte cultures (**Figure 4C**). This family of receptors have essential roles in lipid and cholesterol transport, with LRP1 also implicated in the astrocytic uptake and degradation of amyloid-β [106–108]. The low-density lipoprotein receptors have been shown to interact with SNX17 [109, 110] through their NPxY motifs within their intracellular domains, with SNX17 interacting with the endosomal sorting complex Retriever [111–114]. We validated the loss of LRP1 surface expression biochemically using surface biotinylation, streptavidin affinity capture coupled with quantitative western analysis (**Figure 4D-E**). VPS35 suppression led to a 57% reduction in surface LRP1 levels (p = 0.0013, n = 3, one-way ANOVA) whilst SNX27 suppression led to a 69% reduction in surface LRP1 levels (p = 0.0004, n = 3, one-way ANOVA). SNX3/SNX12 suppression also caused a small decrease in surface LRP1 expression by 23% (p = 0.0449, n = 3, unpaired t-test) We also quantified a decrease in total levels of LRP1 expression (60% reduction after VPS35 suppression (P < 0.0001, n = 3, one-way ANOVA) and a 72% reduction after SNX27 suppression (P < 0.0001, n = 3, one-way ANOVA)) indicating that a loss of SNX27-Retromer results in LRP1 being degraded in the lysosome. This data highlights that LRP1 expression can be affected by SNX27-Retromer in astrocytes.

### Role of SNX27-Retromer in glutamate homeostasis

Comparing across the neuronal and astrocytic proteomic datasets 31 proteins were depleted from the cell surface in both neurons and astrocytes (**Figure 5A, Supplementary Figure 5A**) and 7 proteins were enriched in both cell types following VPS35 suppression (**Figure 5B, Supplementary Figure 5B**). This limited crossover highlights the importance of investigating the effect of Retromer perturbation in different cell types. When comparing our proteomic data across the neuronal and astrocyte cultures of particular interest was the loss of surface expression of the glutamate transporters (SLC1A1/EAAT3, SLC1A2/EAAT2/GLT1 and SLC1A3/EAAT1/GLAST) after SNX27-Retromer suppression. These transporters are essential to remove glutamate from the synaptic cleft, terminating neuronal transmission and thereby preventing aberrant glutamatergic signalling and excitotoxicity [115, 116]. SLC1A1, like SLC1A3, contains a Type I PDZbm in its carboxy-terminal tail indicative of high affinity binding to SNX27 with our data suggesting that these transporters depend on the SNX27-Retromer complex for their trafficking. For SLC1A2 only one of the three isoforms detected in rat contain a PDZbm in its carboxy-terminal tail (isoform b, NCBI Reference Sequence: NP_001030310.2). AlphaFold2 also predicted direct binding between SNX27 and the PDZbm of SLC1A1 and SLC1A3 with an average iPTM score of 0.744 and 0.784 respectively (**Supplementary Figure 5C, Supplementary Tables 4, 7**). In total, 42 neuronal and 22 astrocytic cargoes were predicted by AlphaFold2 to interact with the PDZ of SNX27 through their PDZbm with iPTM scores over 0.6 (**Supplementary Figure 5D, Supplementary Tables 4, 7)**.

**Figure 5.**
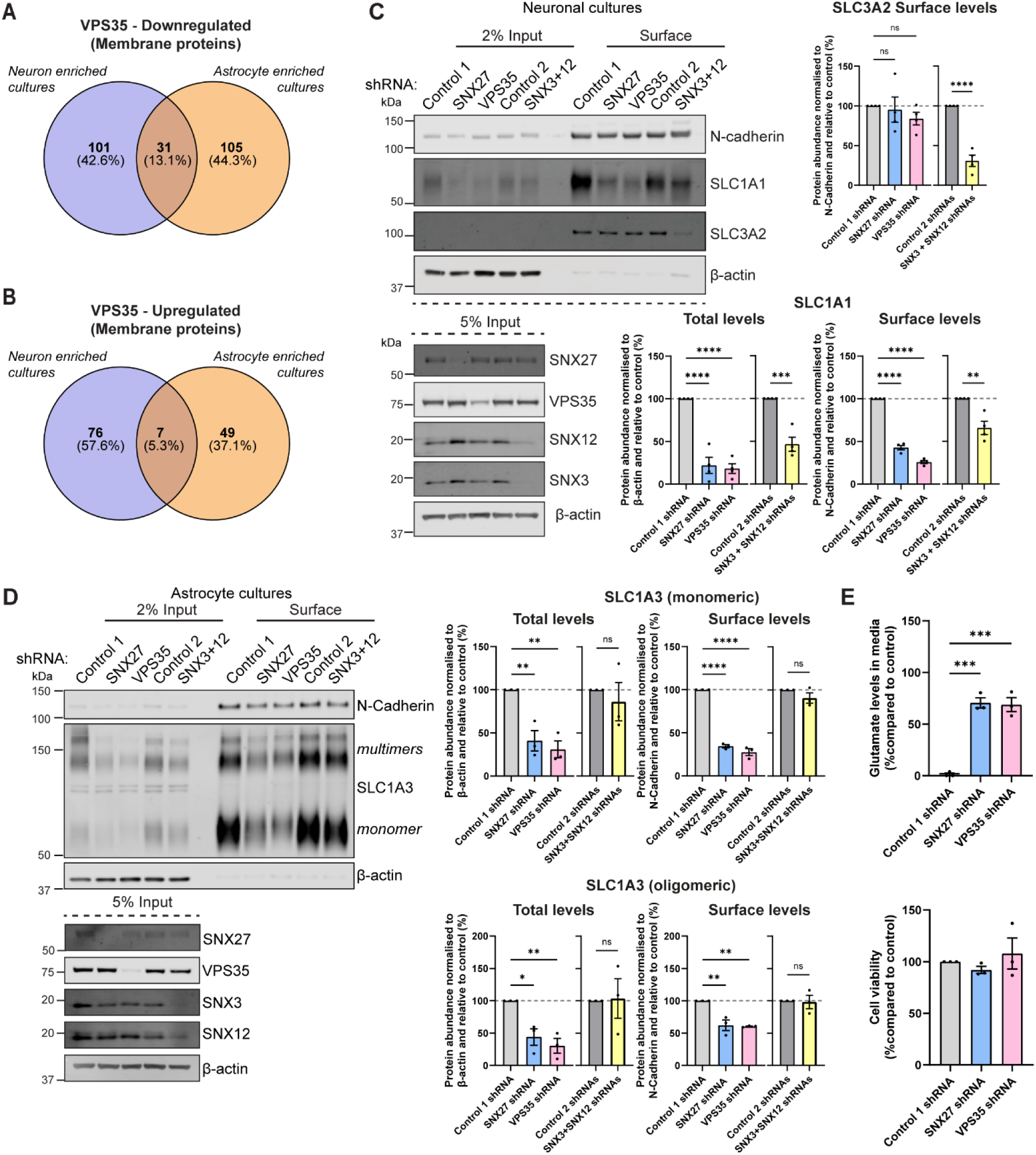
Role of SNX27-Retromer in glutamate homeostasis. **(A-B)** Venn diagrams comparing the number of membrane proteins significantly **(A)** downregulated or **(B)** upregulated at the surface of rat cortical astrocyte and rat cortical neuronal cultures after VPS35 suppression. **(C)** Fluorescence-based western analysis after surface biotinylation and streptavidin agarose capture of membrane proteins of rat cortical neurons transduced with control non-targeting shRNAs (Control 1 and 2) or shRNAs against SNX27, VPS35 or SNX3+SNX12. Endogenous surface and total levels of the glutamate transporter SLC1A1 and SLC3A2, and control N-Cadherin are shown. The level of VPS35, SNX27 and SNX3+SNX12 are also shown with β-actin used as a protein load control. Quantification from western analyses from four independent experiments (n = 4). Surface SLC1A1 and SLC3A2 are normalised to control N-Cadherin whilst total levels are normalised to β-actin. **(D)** Fluorescence-based western analysis after surface biotinylation and streptavidin agarose capture of membrane proteins of rat cortical astrocytes transduced with control non-targeting shRNAs (Control 1 and 2) or shRNAs against SNX27, VPS35 or SNX3+SNX12. Endogenous surface and total levels of glutamate transporter SLC1A3 and control N-Cadherin are shown. The level of VPS35, SNX27 and SNX3+SNX12 are also shown with β-actin used as a protein load control. Quantification from western analyses from three independent experiments (n = 3). Surface SLC1A3 are normalised to control N-Cadherin whilst total levels are normalised to β-actin. Data expressed as a percentage of the relevant control shRNA. **(E)** Glutamate assay measuring the amount of glutamate in the media in rat astrocyte cultures after suppression of SNX27 or VPS35. Cell viability assay showed no significant changes. **(C-E)** SNX27 and VPS35 shRNA data analysed by an one-way ANOVA followed by a post hoc Dunnett’s test compared to Control 1. SNX3+SNX12 shRNA data analysed by an unpaired t-test compared to Control 2. Error bars represent mean ± SEM. ****, p ≤ 0.0001; ***, p ≤ 0.001; **, p ≤ 0.01; *, p ≤ 0.005; ns, not significant.

SLC1A1 is thought to be primarily expressed in neurons particularly at the pre-synapse whilst SLC1A2 and SLC1A3 are thought to be primarily astrocytic [117]. To validate whether SNX27-Retromer suppression alters the surface expression of these glutamate receptors we used surface biotinylation followed by streptavidin pulldowns and western analysis in either shRNA treated astrocyte cultures or neuronal cultures. Unfortunately, we could not find an antibody to reliably detect SLC1A2 in our rat cultures. In the SNX27 and VPS35 suppressed neuronal cultures we found a significant decrease in SLC1A1 surface expression by 57% (p = <0.0001, n = 4, one-way ANOVA) and 74% (p = <0.0001, n = 4, one-way ANOVA) respectively.

We also found a significant decrease in the total levels of SLC1A1 by 78% (p = <0.0001, n = 4, one-way ANOVA) after SNX27 suppression and 82% (p = <0.0001, n = 4, one-way ANOVA) after VPS35 suppression suggesting that the mis-trafficked SLC1A1 is being degraded in the lysosome (**Figure 5C**). SNX3/SNX12 suppression also showed a smaller but significant loss of SLC1A1 at both the surface (34% reduction, p = 0.0043, n = 4, one-way ANOVA) and total level (53% reduction, p = 0.0006, n = 4, unpaired t-test).

In the SNX27 and VPS35 suppressed astrocyte cultures we found that the surface expression of monomeric SLC1A3 was significantly decreased by 66% (p = <0.0001, n = 3, one-way ANOVA) and 73% (p = <0.0001, n = 3, one-way ANOVA), respectively, with the multimeric forms of SLC1A3 also showing a reduction. Total levels of SLC1A3 were also shown to be reduced by 59% (p = 0.0059, n = 3, one-way ANOVA) after SNX27 suppression and 69% (p = 0.0027, n = 3, one-way ANOVA) after VPS35 suppression, again indicating that a loss of SNX27-Retromer results in SLC1A3 being degraded in the lysosome **(Figure 5D)**. SNX3/SNX12 suppression did not significantly affect SLC1A3 expression highlighting that SLC1A3 is a SNX27-Retromer cargo. To test whether this loss of surface glutamate transporters affected glutamate uptake in our astrocyte cultures we added 100 µm glutamate and measured the level of glutamate in the media after 1 hour. We found a significant increase in the media glutamate levels in the SNX27 (69% increase, p = 0.0001) and VPS35 shRNA (68% increase, p = 0.0001) treated astrocyte cultures compared to the control 1 non-targeting shRNA treated cells (**Figure 5E)**, consistent with suppressed astrocytes having a reduced efficiency in glutamate uptake. This loss of uptake was not due to a loss of cell viability (**Figure 5E)**. These data highlight an important role for Retromer in the trafficking of glutamate transporters in both neurons and astrocytes and highlight a pathway, if perturbed, could lead to excitotoxicity and neuronal death.

In addition to its role as an excitatory neurotransmitter, glutamate also plays other important roles in cells including imparting antioxidant effects via the synthesis of reduced glutathione (GSH) [118]. The synthesis of GSH is primarily regulated by system XC^-^, comprised of the single-pass transmembrane protein SLC3A2 (4F2hc), which is thought to act as a chaperone for the membrane localisation and stability of the glutamate/cysteine antiporter SLC7A11 [119]. In our proteomic analysis we identified a reduction in surface levels of SLC3A2, in both cortical neurons and astrocytes, with suppression of SNX3/12 and VPS35 (**Supplementary Table 1 and 6**). SNX3/12 suppression additionally led to the reduction of surface levels of SLC7A11 in both cell types, with VPS35 suppression also leading to a significant reduction in cortical neurons (**Figure 2C**). Surface biotinylation followed by western blotting validated a significant 69% loss of SLC3A2 from the surface of neurons following SNX3/12 suppression (p < 0.0001, n = 4, unpaired t-test) (**Figure 5C**). AlphaFold analysis did not predict a confident interaction of SLC3A2 with SNX3-Retromer across models tested, indicating that the reduction in surface expression of SLC3A2 with SNX3/SNX12 suppression may be an indirect consequence, for example through trafficking of an independent adaptor protein (**Supplementary Table 5**). Previous work also did not find a change in surface recycling of another binding partner of SLC3A2, SLC7A5 (LAT1, system L), in VPS35 knockout HeLa cells, and at the sensitivity of western blotting we did not identify a change in surface SLC3A2 levels with VPS35 suppression (**Figure 5C**), indicating that additional regulatory mechanisms likely contribute to SNX3-Retromer mediated recycling of SLC3A2 heterodimers [120]. Taken together our proteomic analyses highlights several key pathways and proteins involved in glutamate homeostasis that are affected by Retromer, SNX27 and SNX3/SNX12 suppression across both neurons and astrocytes.

## Discussion

Within the context of neurodegenerative disease, research into the endosomal network has primarily focused on neurons, with little consideration given to glial cells including astrocytes. In the present study, we have addressed this knowledge gap. First, we have shown that perturbation of Retromer and the sorting nexin cargo adaptors induce endosomal swelling in astrocytes, highlighting that these cells are also sensitive to changes in endosomal function. Secondly, by identifying multiple proteins across neurons and astrocytes that display perturbed cell surface recycling in response to endosomal dysfunction we have identified the mechanistic basis for the recycling of several key disease associated pathways including the glutamatergic and GABAergic pathways, APP processing and amyloid response, and the lipoprotein receptor family. Collectively, these data expand our appreciation of the neuroprotective role of Retromer and the endosomal network in aging-related neurodegenerative disease.

### Retromer and the glutamatergic and GABAergic pathways

One of the key pathways affected across both neurons and astrocytes after suppression of SNX27-Retromer and SNX3/SNX12-Retromer is the glutamatergic pathway. There are several studies linking Retromer dysfunction to changes in AMPA and NMDA receptor expression and activity in neurons [80–85]. These ionotropic glutamate receptors are essential for excitatory synaptic transmission and synaptic activity, which are important in fundamental central processing. Many of the AMPA and NMDA receptor subunits were significantly reduced in the VPS35 suppressed neuronal cultures but, interestingly, we also found a decrease in surface expression of AMPA receptor subunits in our astrocyte-enriched cultures (with less than 5% of these cells being neurons). AMPA and NMDA receptors have been shown to be expressed in astrocytes with roles in glial excitability and neuronal activity [121, 122]. However, future work is needed to address how Retromer suppression affects astrocytic ionotropic glutamate receptor expression and whether their mis-trafficking has a role in neuronal function and neurodegenerative disease. Moreover, our data has highlighted a role for Retromer in the trafficking of metabotropic glutamate receptors, which are involved in intracellular signalling pathways leading to changes in synaptic function, and the excitatory amino acid transporters which are important for glutamate uptake. We validated that Retromer suppression leads to a loss of surface expression of the neuronal glutamate transporter SLC1A1/EAAT3 and a loss of surface expression of the astrocytic glutamate transporter SLC1A3/EAAT1, along with SLC1A2/EAAT2. SLC1A2 and SLC1A3 are primarily expressed in astrocytes and are important in removing most of the glutamate from the synaptic cleft. Impairment of glutamate uptake leads to overexcitation of glutamate receptors present on the post-synapse leading to elevated calcium influx, and ultimately excitotoxicity and neuronal death [86, 123]. Excitotoxicity has been associated with several neurodegenerative disorders as well as epilepsy [124, 125]. Indeed, knockout of SNX27 in mice leads to severe neuronal death and early lethality [65] and SNX27 mutations in humans have been linked to epileptic seizures [63, 64]. Our data show that surface expression of glutamate transporters SLC1A2 and SLC1A3 are significantly decreased after SNX27-Retromer suppression and that these astrocytic cultures display a reduce ability to efficiently uptake glutamate. We speculate that this will likely result in neuronal over activation, excitotoxicity and neuronal death.

In addition to the glutamatergic pathway, our analyses also highlighted the GABAergic pathway as a key pathway affected after Retromer suppression, with many of the GABA receptors subunits decreased at the neuronal surface. Indeed, disturbances in the glutamatergic and GABAergic systems have been heavily implicated in neurodegenerative pathology [86–88, 123, 126–128], and previous papers have reported an effect of SNX27 depletion on inhibitory synapse function through trafficking of the inhibitory synapse adhesion molecule Neuroligin-2 (NLG2) [77, 129]. Therefore, it is currently unclear whether the reduction in GABA receptors represents an indirect effect via NLG2, a compensatory effect to maintain the excitatory/inhibitory balance upon loss of glutamatergic signalling, or whether Retromer-SNX complexes can directly interact with and traffic GABA receptors through the endosomal system. Experiments are currently underway to explore the complexities of such an integrated network.

### Global analysis of SNX3-Retromer cargo

To our knowledge this is the first global analysis of SNX3/SNX12-Retromer cargo. Although identification of SNX3/SNX12 direct cargo is complicated by the contribution of SNX3/SNX12 to the recruitment of Retromer onto endosomal membranes, AlphaFold modelling predicts an interaction of the cytoplasmic tails of a select group of these proteins with SNX3-Retromer, indicating these are recycled via sequence-dependent cargo sorting. The relatively low number of high confidence predictions for a direct interaction with SNX3-Retromer, however, may indicate that many of the proteins significantly altered in this condition are an indirect consequence of the role of SNX3/SNX12 in Retromer localisation to the endosomal retrieval subdomain. The variability of some predictions across input sequences and version releases, also highlights the lack of clarity of the permitted residues within the SNX3-Retromer cargo binding motif. Alternatively, our *in-silico* modelling is lacking post-translational modifications and membrane interactions, which may be required for the binding of SNX3-Retromer to cargo.

Our proteomic analyses identified a depletion of system XC^-^ (SLC3A2/SLC7A11) on the cell surface upon suppression of SNX3/SNX12 expression. System XC^-^ plays a key role in maintaining redox homeostasis inside cells via the uptake of cystine in exchange for glutamate, a key step in the synthesis of GSH required for reactive oxygen species (ROS)-detoxifying enzymes [119]. Recent work identified oxidation of cysteine residues in VPS35 in response to intracellular ROS as a preventative mechanism against oxidative stress, highlighting Retromer as a modulator of this pathway [130]. SLC7A11 has recently been shown to function on lysosomal membranes, with cystine/glutamate exchange providing a mechanism for preventing lysosomal over-acidification in the presence of high H^+^ levels. Increased SLC7A11 lysosomal levels upon SNX3-Retromer deficiency may therefore contribute to the lysosomal acidification defects observed upon loss of VPS35 expression [131, 132]. SLC7A11 methylation and rare coding variants have also been linked to Parkinson’s Disease risk [132, 133]. Future work is required to determine if SNX3-Retromer plays a direct or indirect role in trafficking of system XC^-^.

### Retromer, APP processing and amyloid-β uptake

Changes in APP processing and the formation of toxic amyloid-β aggregates are a key characteristic of Alzheimer’s disease pathology [89]. Retromer, SNX3 and SNX12 have all been shown to affect APP processing and trafficking [55, 73, 103–105, 134], with this study further highlighting a significant role. Previous studies have demonstrated that VPS35 deficiency specifically increases β-cleavage of APP, which is primarily thought to be driven by the reduced trafficking of APP away from endosomes, resulting in increased exposure to BACE1 cleavage [135]. However, the α-secretase cleavage of APP by ADAM10 has also been shown to be inversely related to BACE1 mediated β-secretase cleavage [101, 102, 136]. In our proteomic data in neurons, we found an increase in APP and BACE1 at the surface and a decrease in ADAM10 with SNX3/SNX12-Retromer suppression. AlphaFold modelling additionally predicted a direct interaction of ADAM10 with SNX3-Retromer, with a binding motif consistent with previously published SNX3-Retromer cargo [66]. Further studies are required to determine if the altered surface levels of these secretases contribute to the increased β-cleavage of APP upon VPS35 suppression. SNX27 has been shown to interact with Presenillin-1, regulating γ-secretase activity, with SNX27 depletion leading to increased Aβ production [137]. Furthermore, SNX27 and Retromer interact with the sortilin-related receptor SORLA, which traffics APP and limits its amyloidogenic proteolysis [12, 138–140]. Interestingly, in the astrocyte proteomics SORLA was significantly reduced at the surface after suppression of VPS35, SNX27 and SNX3/SNX12, which may also affect APP processing. As has been shown previously, VPS35 deficiency also leads to a similar increase in APP-related proteins APLP1 and APLP2 at the cell surface, which are thought to be processed by secretases in the same way as APP [79, 93, 141].

We hypothesise that the increase in APP and related proteins on the surface of both neurons and astrocytes is due to an increase in lysosomal exocytosis, based on our proteomic data also showing a significant increase of lysosomal proteins at the cell surface, as well as previous data illustrating an essential role for Retromer in lysosomal health [79]. Perturbation of the lysosomal system and an increase in lysosomal exocytosis are increasingly being associated with neurodegenerative disease [79, 142–146]. In addition, our study has highlighted proteins involved in amyloid-β uptake as being significantly affected after Retromer suppression. SNX27-Retromer suppression led to a significant loss of surface levels of lipoprotein receptor-related protein 1 (LRP1) in astrocytes, which has been implicated in the uptake and degradation of amyloid-β and tau [107, 147]. Impaired clearance of amyloid-β leads to its aggregation and subsequent plaque formation, both of which are hallmarks of Alzheimer’s pathology. Several members of the low-density lipoprotein receptor family were affected by SNX27-Retromer suppression in astrocytes, with these receptors having essential roles in lipid and cholesterol transport. The low-density lipoprotein receptors contain NPxY motifs and have been shown to depend on the SNX17-Retriever endosomal sorting complex for their trafficking [109, 110]. SNX17-Retriever assembles into a larger complex, Commander, with perturbation of this complex implicated in neurodevelopmental disorders [111–114, 148–150]. Assessing the interplay between SNX17-Retriever and SNX27-Retromer in regulating the neuronal and astrocytic cell surface recycling of lipoprotein receptors will be an important future direction.

### Retromer’s role across different cell types in the brain

Very few studies have looked at the role of Retromer or the SNXs in non-neuronal cells of the brain. However, glial cells including astrocytes, microglia and oligodendrocytes, are essential for brain homeostasis and are increasingly being associated with neurodegenerative disease [151]. Retromer depletion in neurons has been shown to increase gliosis [82, 131], and lead to dysmorphic microglial morphology in mice, phenotypic of the microglial morphology observed in AD patients [82]. VPS35 depletion in microglia leads to microglial activation and a loss of neurogenesis in mice [152], and accelerates AD pathology in the 5XFAD mouse model [153]. Retromer has also been implicated in the trafficking of the microglia receptor Triggering receptor expressed on myeloid cells 2 (TREM2), with mutations in TREM2 associated with an increased risk for developing AD [154, 155]. Indeed, a mutation in SNX27 (R198W) associated with intellectual disability has been shown to effect astrocyte metabolism, leading to synaptic and learning deficits in mice [156]. Moreover, SNX27 traffics the G protein-coupled receptor 17 (GPR17), which plays crucial roles in myelination in oligodendrocytes [157], and has been implicated in Aquaporin 4 (AQP4) trafficking with a loss of AQP4 in leading to ventriculomegaly [158]. These studies highlight a key role for Retromer in glial function, with our data expanding the functional role of Retromer in astrocytes.

### Conclusion

Accumulating evidence has highlighted Retromer as a major neuroprotective component across an array of neurodegenerative diseases. Our study provides new insight and increases our understanding of the cargoes that depend on Retromer and its cargo adaptors SNX27 and SNX3/SNX12 for their surface expression in both neurons and astrocytes. Our work identifies new avenues of research and the potential for additional therapeutic targets in the treatment of neurodegenerative disease.

## Methods

### Antibodies

| Antibody | Source | Identifier | Additional information |
| --- | --- | --- | --- |
| GFAP | Dako | Z0334 | IF (1:500) |
| GFAP | Antibodies.com | A85307 | IF (1:500) |
| SLC1A3/GLAST | Proteintech | 20785-1-AP | IF (1:400) |
| IBA1 | Antibodies.com | A82670 | IF (1:200) |
| S100β | Sigma-Aldrich | SAB4200671) | IF (1:200) |
| TUJ1/ βIII-tubulin | Sigma-Aldrich | T8578 | IF (1:200) |
| SNX6 | Santa Cruz Biotechnology | sc-365965 | IF (1:200) |
| VPS35 | Abcam | Ab97545 | IF (1:200) |
| EEA1 | Cell Signaling Technology | 3288 | IF (1:200) |
| VPS35 | Abcam | Ab157220 | WB (1:2000) |
| SNX27 | A kind gift from Dr Martin Playford |  | WB (1:500) |
| SNX3 | Proteintech | 10772-1-AP | WB (1:1000) |
| SNX12 | Santa Cruz Biotechnology | Sc-101308 | WB (1:500) |
| $\beta$ -actin | Sigma-Aldrich | A1978 | WB (1:10,000) |
| N-cadherin | Cell Signaling Technology | 14215S | WB (1:1000) |
| SLC1A3/GLAST | Proteintech | 20785-1-AP | WB (1:1000) |
| SLC1A1 | Cell Signaling Technology | 14501 | WB (1:1000) |
| APP | Abcam | ab32136 | WB (1:1000) |
| LAMP1 | Cell Signaling Technology | 3243 | WB (1:1000) |
| SLC3A2 | Cell Signaling Technology | 13180 | WB (1:1000) |
| SLC4A7 | Invitrogen | PA5-25563 | WB (1:1000) |
| LRP1 | Abcam | Ab92544 | WB (1:1000) |

### Cell culture

All cells were cultured in a humidified incubator at 37°C and 5% CO_2_. HEK293T cells (American Type Culture Collection (ATCC)), for lentiviral production, were maintained in DMEM (D5796; Sigma-Aldrich) supplemented with 10% foetal bovine serum (F7524; Sigma-Aldrich) and 1% Penicillin/Streptomycin (15070063; Thermo Fisher Scientific). Rat primary cultures were prepared in accordance with the UK Animal (Scientific Procedures) Act 1986. Primary cultures were prepared from embryonic day E18 Wistar rat brains as previously described [159]. Briefly, for neuronal cultures dissociated cortical cells were grown in six well dishes (500,000 cells/well), coated with poly-L-lysine (0.5 mg/ml; P2636; Sigma-Aldrich), in 2 ml plating medium (Neurobasal medium (21103–049; Gibco) supplemented with 5% horse serum (H1270; Sigma-Aldrich), 2% B27 (17504–044; Gibco), 1% Penicillin/Streptomycin, and 1% Glutamax (35050–038; Gibco)). The media was exchanged for 2 ml feeding medium 2 hr after plating (Neurobasal medium (21103–049; Gibco), 2% B27 (17504–044; Gibco), 0.4% Penicillin/Streptomycin, and 0.4% Glutamax (35050–038; Gibco)), with the cells fed an additional 1 ml of feeding medium 7 days after plating.

For astrocyte cultures, dissociated cortical cells were grown in a T75 flask (10-15 million cells/flask), coated with poly-D-lysine (P0899, Sigma-Aldrich), in 20 ml of astrocyte media (DMEM (D5796; Sigma-Aldrich) supplemented with 10% foetal bovine serum (F7524; Sigma-Aldrich) and 1% Penicillin/Streptomycin (15070063; Thermo Fisher Scientific)). After 7 days (changing the media every 2-3 days) the astrocytes were separated from the microglia by being shaken on an orbital shaker at 180 rpm for 30 min. The media (containing the microglia cells) was removed and a fresh 20 ml of astrocyte media added. The cells were shaken again at 300 rpm for 6 h to separate the oligodendrocyte precursor cells, the media removed, and the cells washed in PBS and trypsinised in 1X Trypsin-EDTA diluted in PBS (T4174; Sigma-Aldrich). Once the astrocyte cells were detached, astrocyte media was added and the cells transferred to a 15ml falcon and centrifuged at 1200 x rpm for 5 min. The media was aspirated, cells resuspended in astrocyte media and split into 2x T75 flasks. After a further 7 days (changing media every 2-3 days) cells were split into 2x T175 flasks and left to grow for a further 6-7 days before being plated into 6-well plates for experiments (200,000 cells/well).

### shRNA lentiviral production

shRNAs driven by a H1 promoter were generated for the knockdown of rat SNX27 (target sequence 5’-aagaacagcaccacagaccaa-3’) [59, 77], rat VPS35 (target sequence 5’-aacagtggagatattcaataa-3’), rat SNX3 (target sequence 5’-gaacgttgtcttcacatgt-3’) or the non-targeting control shRNA (non-targeting sequence 5’-aattctccgaacgtgtcac-3’). Oligonucleotides were cloned into a modified pXLG3-GFP viral vector containing a Woodchuck Hepatitis Virus Posttranscriptional Regulatory Element (WPRE) to increase the GFP expression. For the rat SNX12 shRNA (target sequence 5’-aacttcctggagatagacatc-3’), as well as the non-targeting control shRNA (non-targeting sequence 5’-aattctccgaacgtgtcac-3’), the oligonucleotides were cloned into a modified pXLG3-mcherry-WPRE viral vector. The viral vectors were co-transfected into a 15 cm dish of HEK293T cells with the helper plasmids psPax2 and pMDG2 using PEI (23966, Polysciences). The viruses were harvested 48 hr after transfection, spun down at 4000 rpm for 10 min at room temperature and filtered through 0.45 μm filter caps before being stored at −70°C. Neurons were transduced with shRNA viruses on DIV12 and left for 7 days before analysis. Astrocytes were plated on DIV18, transduced on DIV19 and left for 7 days before analysis. To control for the SNX3 and SNX12 double suppression, non-targeting controls co-expressing both GFP and mCherry were used concurrently (Control 2).

### Surface biotinylations

All solutions were pre-chilled to 4°C and all steps were carried out on ice to prevent internalisation. Fresh membrane impermeable Sulpho NHS-SS-Biotin (21331; Thermo Fisher Scientific) was dissolved in Tyrode’s buffer (25 mM HEPES, 119 mM NaCl, 2.5 mM KCl, 2 mM CaCl_2_, 2 mM MgCl_2_, 30 mM glucose, pH 7.4) at a final concentration of 0.2 mg/ml. Cells were washed twice in Tyrode’s buffer before being incubated with biotin for 15 mins at 4°C. To remove excess biotin the cells were washed in Tyrode’s buffer before being quenched twice in Tyrode’s quenching buffer (25mM HEPES, 19 mM NaCl, 2.5 mM KCl, 2mM CaCl_2_, 2 mM MgCl_2_, 30 mM glucose, 100 mM NH_4_Cl, pH 7.4) for 1 min at 4°C. The cells were washed a final time in Tyrode’s buffer before being lysed in 2% Triton-X-100 (X100; Sigma) plus 0.1% SDS (15553027; Thermo Fisher Scientific) with protease inhibitor cocktail tablets (A32955; Thermo Fisher Scientific) and phosphatase inhibitor tablets (A32957; Thermo Fisher Scientific) in PBS. The lysate was centrifuged at 14, 000 rpm for 10 min at 4°C and transferred to a fresh microcentrifuge tube. A BCA assay (23225; Thermo Fisher Scientific) was carried out to determine protein concentration. An aliquot of the lysate was retained to represent the whole cell fraction and equal protein amounts of lysate were incubated with prewashed (in lysis buffer) streptavidin beads (17-5113-01; GE Healthcare) for 1 hr at 4°C. The biotin precipitated beads were washed once in wash buffer 1 (1% Triton-X-100 (X100; Sigma-Aldrich) in PBS), twice in wash buffer 2 (1% Triton-X-100 (X100; Sigma-Aldrich), 1 M NaCl in PBS) and once in PBS. For proteomic samples 30 μl PBS was added to the biotin precipitated beads and the samples flash frozen in liquid nitrogen before being stored at −80°C before analysis. For western analysis beads were resuspended in a 2x loading buffer, containing 2.5 % β-mercaptoethanol (M3148; Sigma-Aldrich) and all samples were denatured at 95 °C for 5 min.

### Western blot analysis

Proteins were resolved on NuPAGE 4–12% precast Bis-Tris gels (NP0336BOX; Invitrogen) and then transferred onto methanol activated polyvinylidene fluoride (PVDF) membranes (10600029; GE Healthcare), before being blocked in 5% milk in Tris-buffered saline (150 mM NaCl, 10 mM tris pH 7.5), plus 0.1% Tween (TBS-T) for 1 h at room temperature. The membranes were then incubated with the appropriate primary antibodies’ overnight at 4°C. The membrane was washed in TBS-T before being incubated with Alexa Fluor secondary antibodies (1: 10 000 of mouse 680 or rabbit 800; Invitrogen). After washing in TBS-T the protein bands were visualised using an Odyssey infrared scanning system (LI-COR Odyssey CLX). Quantification of band intensities was performed in Image Studio Lite software.

### Proteomic analysis

#### TMT Labelling and High pH reversed-phase chromatography

Streptavidin-isolated samples were reduced (10mM TCEP, 55°C for 1h), alkylated (18.75mM iodoacetamide, room temperature for 30min) and then digested from the beads with trypsin (1.25 µg trypsin; 37°C, overnight). The resulting peptides were labelled with Tandem Mass Tag (TMT) six plex reagents according to the manufacturer’s protocol (Thermo Fisher Scientific, Loughborough, LE11 5RG, UK) and the labelled samples pooled.

The pooled sample was desalted using a SepPak cartridge according to the manufacturer’s instructions (Waters, Milford, Massachusetts, USA). Eluate from the SepPak cartridge was evaporated to dryness and resuspended in buffer A (20 mM ammonium hydroxide, pH 10) prior to fractionation by high pH reversed-phase chromatography using an Ultimate 3000 liquid chromatography system (Thermo Fisher Scientific). In brief, the sample was loaded onto an XBridge BEH C18 Column (130Å, 3.5 µm, 2.1 mm X 150 mm, Waters, UK) in buffer A and peptides eluted with an increasing gradient of buffer B (20 mM Ammonium Hydroxide in acetonitrile, pH 10) from 0-95% over 60 minutes. The resulting fractions (concatenated into 5 in total) were evaporated to dryness and resuspended in 1% formic acid prior to analysis by nano-LC MSMS using an Orbitrap Fusion Lumos mass spectrometer (Thermo Fisher Scientific).

#### Nano-LC Mass Spectrometry

High pH RP fractions were further fractionated using an Ultimate 3000 nano-LC system in line with an Orbitrap Fusion Lumos mass spectrometer (Thermo Fisher Scientific). In brief, peptides in 1% (vol/vol) formic acid were injected onto an Acclaim PepMap C18 nano-trap column (Thermo Fisher Scientific). After washing with 0.5% (vol/vol) acetonitrile 0.1% (vol/vol) formic acid peptides were resolved on a 250 mm × 75 μm Acclaim PepMap C18 reverse phase analytical column (Thermo Fisher Scientific) over a 150 min organic gradient, using 7 gradient segments (1-6% solvent B over 1min., 6-15% B over 58min., 15-32%B over 58min., 32-40%B over 5min., 40-90%B over 1min., held at 90%B for 6min and then reduced to 1%B over 1min) with a flow rate of 300 nl min^−1^. Solvent A was 0.1% formic acid, and Solvent B was aqueous 80% acetonitrile in 0.1% formic acid. Peptides were ionized by nano-electrospray ionization at 2.0kV using a stainless-steel emitter with an internal diameter of 30 μm (Thermo Fisher Scientific) and a capillary temperature of 300°C.

All spectra were acquired using an Orbitrap Fusion Lumos mass spectrometer controlled by Xcalibur 3.0 software (Thermo Fisher Scientific) and operated in data-dependent acquisition mode using an SPS-MS3 workflow. FTMS1 spectra were collected at a resolution of 120 000, with an automatic gain control (AGC) target of 400 000 and a max injection time of 100ms. Precursors were filtered with an intensity threshold of 5000, according to charge state (to include charge states 2-7) and with monoisotopic peak determination set to Peptide. Previously interrogated precursors were excluded using a dynamic window (60s +/-10ppm). The MS2 precursors were isolated with a quadrupole isolation window of 0.7m/z. ITMS2 spectra were collected with an AGC target of 10 000, max injection time of 70ms and CID collision energy of 35%.

For FTMS3 analysis, the Orbitrap was operated at 30 000 resolution with an AGC target of 50 000 and a max injection time of 105ms. Precursors were fragmented by high energy collision dissociation (HCD) at a normalised collision energy of 60% to ensure maximal TMT reporter ion yield. Synchronous Precursor Selection (SPS) was enabled to include up to 10 MS2 fragment ions in the FTMS3 scan.

### Immunofluorescence staining

Neurons were grown on 22 mm glass coverslips coated with poly-L-lysine (1mg/ml; P2636; Merck) and fixed in 4% paraformaldehyde in PBS for 15 min at room temperature before being quenched in 100 mM glycine and washed in PBS. Neurons were permeabilised in 0.1% Triton X-100 (X100; Sigma) in PBS for 5 min at room temperature. Neurons were blocked with 2% BSA (A9647; Sigma-Aldrich) in PBS for 10 min at room temperature, followed by incubation for 1 h at room temperature with the indicated primary antibodies diluted in 0.1 % BSA in PBS. Astrocytes were grown on 13 mm glass coverslips coated with 1X geltrex (A14133-02; Thermo Fisher Scientific) and laminin (1:100 dilution; 23017-015; Thermo Fisher Scientific) and fixed in 4% paraformaldehyde (28908; Thermo Fisher Scientific) in PBS for 15 min at room temperature. To remove fixative, cells were washed three times in PBS before permeabilisation in 0.1% Triton X-100 (X100; Sigma) in PBS for 5 min at room temperature. Cells were blocked with 1% BSA (A9647; Sigma-Aldrich) in PBS for 10 min at room temperature, followed by incubation for 1 h at room temperature with the indicated primary antibodies diluted in 0.1 % BSA in PBS. After primary antibody detection, coverslips were washed three times in PBS before incubation with the appropriate Alexa Fluor conjugated secondary antibodies (488, 568 and 647; Invitrogen) and DAPI (D9542; Sigma-Aldrich) for 1 hr at room temperature. The coverslips were washed a final three times in PBS before being mounted on glass microscope slides using Fluoromount-G (00–4958–02; eBioscience).

### Image acquisition and analysis

Z-stack images were captured using a confocal laser-scanning microscope (SP8 Leica Biosystems) attached to an inverted epifluorescence microscope (DMi8 Leica Biosystems). A 63X, NA 1.4, oil immersion objective (Plan Apochromat BL; Leica Biosystems) at 1X zoom with the standard Leica LAS system acquisition software and detector were used. All settings were kept the same within experiments. Fiji ImageJ software (NIH) was used to process all images.

### Glutamate uptake assay

To assess glutamate uptake by astrocytes, we used a glutamate assay kit (ab83389, Abcam). Briefly, rat cortical astrocytes were plated in 6-well dishes. The next day, cells were transduced with lentiviruses, by addition of viral supernatant to the media. 7 days later, cells were incubated with 100 µM glutamate in DMEM containing 10% FBS, 1% Penicillin/Streptomycin and 1% glutamine for 1 hour. The media was removed and the amount of glutamate present measured in duplicate according to the manufacturer’s instructions using a plate reader.

### Cell viability assay

To assess astrocyte viability, astrocytes were plated and virally transduced as above, before replacement of media with growth media containing 10% MTS Assay Reagent (ab197010; Abcam). Cells were then replaced in the incubator for 1 hour for MTS conversion to take place. Viable cell proportion was then determined by measurement of absorbance at 490nm on a plate reader.

### AlphaFold modelling

The top 25 most depleted proteins by logFC (P < 0.05) upon SNX3/SNX12 suppression, with annotated cytoplasmic tail sequences in Uniprot, were selected for analysis. AlphaFold2 modelling was performed using ColabFold multimer v3 with 12 recycles. Input sequence includes the primary isoform listed for *Rattus norvegicus* SNX3 (Q5U211), VPS26A (Q6AY86), VPS35 (Q96QK1) and the cargoes. For each AF run, five ranked models were generated. Model quality and interface confidence were assessed using the interfacial iPTM scores, pDockQ2 score calculated based on ipSAE [160–162].

For AlphaFold3 (AF3) analysis, the AF3 server (https://alphafoldserver.com/) available on the Google platform was used using default setting. The top 25 most depleted proteins mentioned above was analysed together with Retromer and SNX3. The averaged chain-pair iPTM score between VPS26 and each cargo was calculated using the custom script, which it extracted the chain-pair iPTM score from all five models and computed their averaged. To identify cargoes that show confident binding at the VPS26 - SNX3 interface, the previous published crystal structure of VPS26 – VPS35 – SNX3 – DMT1-II (PDB ID: 5F0L) was used as the reference. All five models from each AF3 runs were superimposed and manually inspected using PyMOL. Cargoes cytoplasmic tails showing overlap at the VPS26 – SNX3 interface were recorded. A score of 5 indicates that all five models from a single AF3 run exhibit the same motif binding to the VPS26 – SNX3 interface.

For SNX27, all models mentioned in the manuscript were generated using AlphaFold2 Multimer in batch mode implemented in the Colabfold interface. Default ColabFold settings were used to construct multiple sequence alignments using MMseqs2. For all modelling experiments, we assessed (i) the prediction confidence measures (pLDDT and interfacial iPTM scores), (ii) the plots of the predicted alignment errors (PAE) and (iii) backbone alignments of the final structures. The input sequence included the primary isoform of *Rattus norvegicus* SNX27 (Q8K4V4) and the last 10 amino acids of depleted proteins upon SNX27 suppression. The average iPTM score was calculated from the individual scores of the five generated models in each run.

### Experimental Design and Statistical Rationale

#### Proteomic Data Analysis

The mass spectrometry proteomics data have been deposited to the ProteomeXchange Consortium via the PRIDE (163) partner repository with the dataset identifier PXD078277. Proteomic data were from three independent rat neuronal cortical cultures and three independent rat astrocyte cultures. The raw data files were processed and quantified using Proteome Discoverer software v2.4 (Thermo Fisher Scientific) and searched against the UniProt Rat (neuronal proteomics: downloaded February 2023, 47939 entries; astrocyte proteomics: downloaded February 2024, 47923 entries) using the SEQUEST HT algorithm. Peptide precursor mass tolerance was set at 10ppm, and MS/MS tolerance was set at 0.6Da. Search criteria included oxidation of methionine (+15.995Da), acetylation of the protein N-terminus (+42.011Da), methionine loss from the protein N-terminus (−131.04Da) and methionine loss plus acetylation of the protein N-terminus (−89.03Da) as variable modifications and carbamidomethylation of cysteine (+57.0214) and the addition of the TMT mass tag (+229.163Da) to peptide N-termini and lysine as fixed modifications. Searches were performed with full tryptic digestion and a maximum of 2 missed cleavages were allowed. The reverse database search option was enabled, and all data was filtered to satisfy false discovery rate (FDR) of 5%.

#### Proteomic statistical analysis

The MS data were searched against the Uniprot Rat database as described above, with additional annotation information on 2024-04-03 (Astrocyte) and 2023-08-25 (Neuronal). Protein groupings were determined by PD2.4, however, the master protein selection was improved with an in-house script. The script first searches Uniprot for the current status of all protein accessions and updates redirected or obsolete accessions. The script further takes the candidate master proteins for each group, and uses current Uniprot review and annotation status to select the best annotated protein as master protein without loss of identification or quantification quality.

The protein abundances for each sample were normalised such that all samples had an equal total protein abundance, then both raw and normalised abundances were Log2 transformed to bring them closer to a normal distribution. The data were processed and statistically analysed using normalised abundances. Univariate paired t-tests were performed for each comparison of interest. For all comparisons, the p-value was adjusted using the Benjamini-Hochberg FDR method.

Membrane proteins were identified from Uniprot annotations (type I-type IV and multi-pass). Gene ontology analysis was performed using ClusterProfiler v4.14.6 [163, 164]. The *simplify* function was utilised to remove redundant terms with a similarity cut off of 0.7 and fold enrichment was calculated as geneRatio/bgRatio. The top 20 terms by smallest P-value were visualised using ggplot2 v3.5.2 (R v4.4.0) [165]. For visualisation of proteomics data, volcano plots were generated using the R package EnhancedVolcano [166] and dot plots were generated using ggplot2 v3.5.2 (R v.4.4.0).

#### Western blot data

All statistical analysis was performed on data from a minimum of 3 independent experimental repeats. Total proteins levels were normalised to the protein loading control (β-actin) whilst surface protein levels were normalised to the control protein N-cadherin. The protein levels were then normalised to the relevant control condition (control 1 or 2 shRNAs) and expressed as a percentage. Data were either analysed by a one-way ANOVA followed by a Dunnett’s multiple comparison’s test or an unpaired t-test.

### Glutamate assay and cell viability data

The amount of glutamate present in the media was measured from 2 technical repeats/condition with 2 readings per sample across three independent experiments. The data were normalised to media only conditions (+/- glutamate) and expressed as a percentage. Data were analysed by a one-way ANOVA followed by a Dunnett’s multiple comparison’s test. Cell viability was measured from 2 technical repeats/condition with 4 readings per sample across three independent experiments. The data was normalised to the control shRNA condition and expressed as a percentage. All data were analysed by a one-way ANOVA followed by a Dunnett’s multiple comparison’s test.

### Immunofluorescence staining data

#### Cell type marker staining of astrocyte-enriched cultures

DAPI was used to count the number of cells using Image J across 10 images / experiment (>400 cells/experiment) across 3 independent experimental repeats (30 images in total). The number of GFAP positive, IBAI positive and TUJ1/βIII-tubulin positive cells were counted as well as the number of cells expressing no markers, normalised to the total number of DAPI cells and expressed as a percentage for each image. The mean number of cells from each image and the average mean across the 3 independent experimental repeats were plotted.

#### Endosomal marker EEA1 staining of astrocyte-enriched cultures

Astrocytes and endosomes were identified using the Image J plugin Modular Image Analysis. Any cells that did not express S100β or the shRNA were discounted and the EEA1 Mean Area (µm2) calculated (between 25 – 82 cells analysed per condition/experiment). Statistical analysis was performed on the mean data from 3 independent experimental repeats. Data were normalised to the relevant control condition (control 1 or 2 shRNAs) and expressed as a percentage. SNX27 and VPS35 shRNA data analysed by a one-way ANOVA followed by a post hoc Dunnett’s test compared to Control 1. SNX3/SNX12 shRNA data analysed by an unpaired t-test compared to Control 2.

#### Co-localisation analysis

Co-localisation of SNX6 and VPS35 signals in neuronal cultures was calculated as Pearson’s correlation coefficient using Volocity (v6.3). Cell bodies were manually identified and thresholded for background intensities. Data was analysed by a one-way ANOVA followed by a post hoc Tukey’s test.

#### Statistical analysis of biochemical and cell biological data

All graphs were prepared in GraphPad Prism 10. Individual datapoints represent independent experimental repeats. SNX27 and VPS35 shRNA data analysed by an one-way ANOVA followed by a post hoc Dunnett’s test compared to Control 1. SNX3/SNX12 shRNA data analysed by an unpaired t-test compared to Control 2. Graphs are plotted representing the mean value ± the standard error of the mean (SEM) for each experimental condition. *n* represents the number of independent experimental repeats. In all graphs ****, p ≤ 0.0001; ***, p ≤ 0.001; **, p ≤ 0.01; ***, *, p ≤ 0.05; ns, not significant.

## Acknowledgments

Kirsty McMillan is supported by an Academy of Medical Sciences Springboard Award (SBF0010\1022), a Royal Society Research grant (RG\R1\251280) and a Tenure-Track Fellowship from the University of Liverpool. Work in the Cullen lab is supported by the Wellcome Trust (220260/Z/20/Z and 319367/Z/24/Z), the Medical Research Council (MR/L007363/1 and MR/P018807/1), the Lister Institute of Preventive Medicine, and the award of a Royal Society Research Professorship (RSRP/R1/211004 and RSRPR26/1003). Brett Collins is supported by an Australian National Health and Medical Research Council (NHMRC) Investigator Grant (APP2016410) an Australian Research Council (ARC) Discovery Project Grant (DP240101315) and NHMRC Centre of Research Excellence (APP2035494). Michael Healy is supported by an NHMRC Investigator Grant (APP2042760).

## Author contributions

**Emma Jones:** Conceptualization, Methodology, Investigation, Formal analysis, Writing - original draft, Writing – review and editing. **Helena Adams:** Investigation, Formal analysis, Writing – review and editing. **Kai-en Chen:** Investigation, Formal analysis, Writing – review and editing. **Fabeeha Maroof:** Investigation, Formal analysis, Writing – review and editing. **Therese M. Ibbotson:** Investigation, Formal analysis, Writing – review and editing. **Yasuko Nakamura:** Investigation, Writing – review and editing. **Paul J. Banks:** Investigation, Supervision, Writing – review and editing. **Michael D. Healy:** Investigation, Formal analysis, Writing – review and editing. **Philip A. Lewis:** Formal analysis, Writing – review and editing. **Kate J. Heesom**: Investigation, Formal analysis, Writing – review and editing. **Brett M. Collin:** Conceptualization, Funding acquisition, Supervision, Writing – review and editing**. Kevin A. Wilkinson:** Methodology, Investigation, Formal analysis, Supervision, Writing – review and editing. **Peter J. Cullen:** Conceptualization, Supervision, Funding acquisition, Writing - original draft, Writing – review and editing. **Kirsty J. McMillan:** Conceptualization, Methodology, Investigation, Formal analysis, Supervision, Funding acquisition, Writing - original draft, Writing – review and editing.

## Competing Interest Statement

The authors declare that they have no known competing financial interests or personal relationships that could have appeared to influence the work reported in this paper.

## Supplementary Figures

**Supplementary Figure 1.**
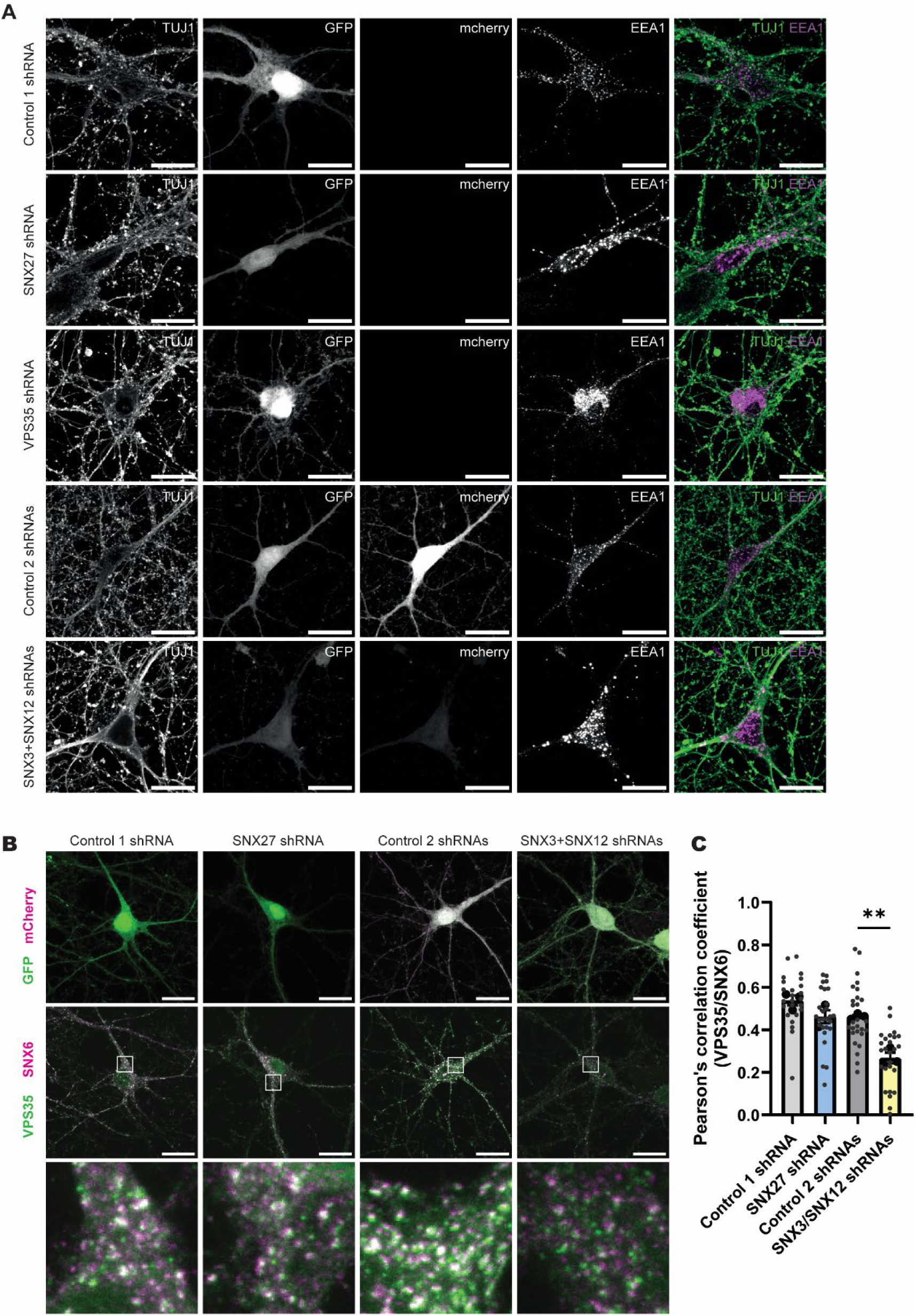
**(A)** Immunofluorescence images of DIV19 rat cortical neurons transduced with constructs co-expressing fluorescent proteins and either control non-targeting shRNAs (Control 1 and Control 2) or shRNAs targeting SNX27, VPS35 or SNX3+12. Neurons were labelled with TUJ1 (pseudo coloured green in merge) and early endosomes were labelled with EEA1 (pseudo coloured pink in merge). Scale bars, 20 μm. (**B**) Immunofluorescent images of DIV19 rat cortical neurons transduced with non-targeting control shRNAs (Control 1 and Control 2) or shRNAs targeting SNX27 or SNX3+SNX12, co-expressing fluorescent proteins GFP and mCherry (top panel: GFP green, mCherry pseudo coloured magenta), co-labelled for endogenous VPS35 with SNX6 as a marker for early endosomal recycling subdomains (bottom panel: VPS35 pseudo coloured green, SNX6 pseudo coloured magenta). Scale bars, 20 μm. (**C**) Pearson’s correlation coefficient of VPS35 and SNX6 localisation, as shown in *(B)*, across three independent experiments (n = 3, 24-27 cells per condition). Data analysed using one-way ANOVA with Tukey’s post-hoc test. Error bars represent mean ± SEM; **, p ≤ 0.01.

**Supplementary Figure 2.**
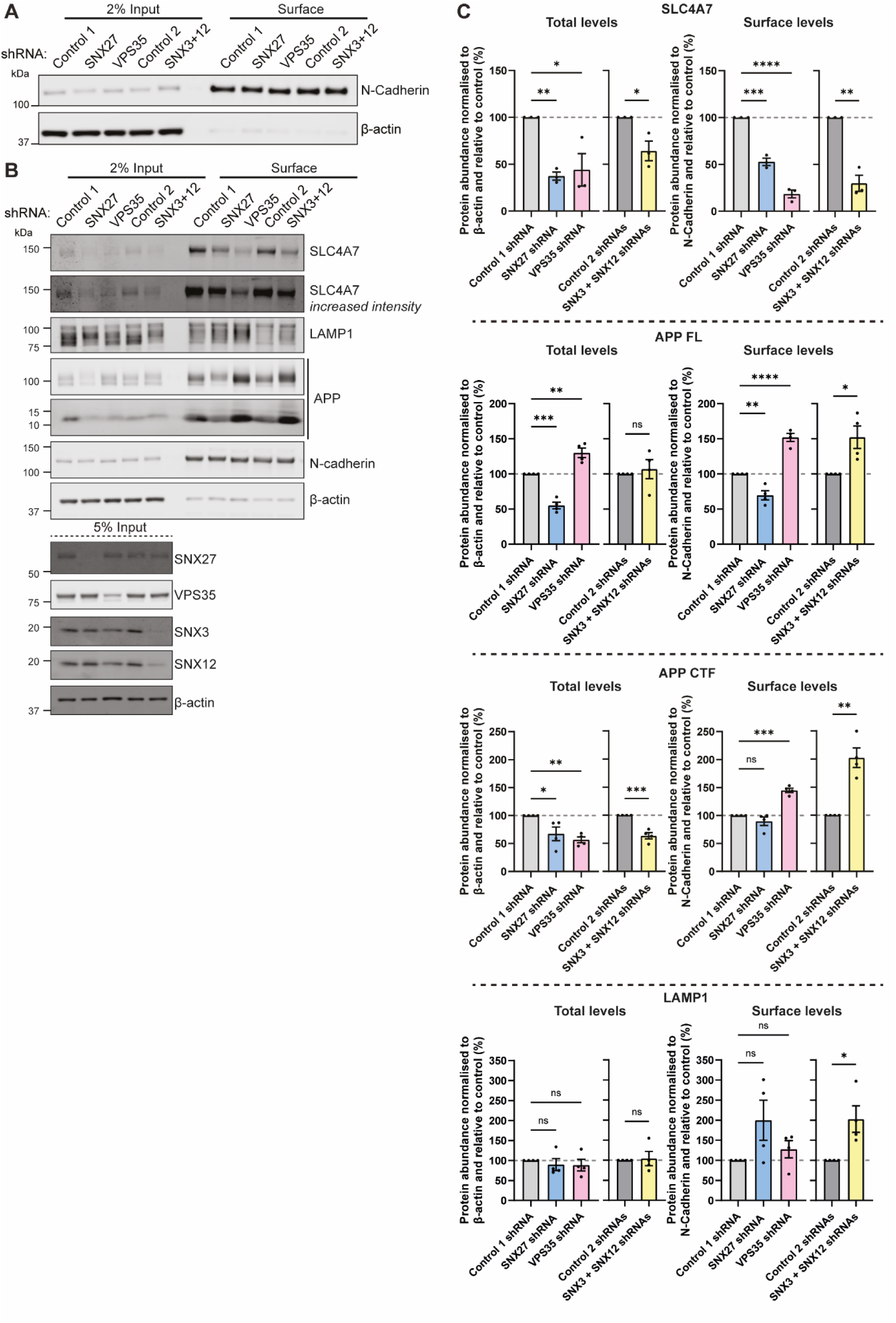
(**A-B**) Fluorescence-based western analysis after surface biotinylation and streptavidin agarose capture of membrane proteins of DIV19 rat cortical neurons transduced with control non-targeting shRNAs (Control 1 and 2) or shRNAs against SNX27, VPS35 or SNX3+SNX12. Endogenous surface and total levels of **(A)** N-cadherin and β-actin and (**B**) SLC4A7, LAMP1, APP (Full length (FL) and C-terminal tail (CTF)), N-cadherin and β-actin. The expression level of VPS35, SNX27 and SNX3+SNX12 are also shown with β-actin used as a protein load control. (**C**) Quantification from western analyses of *(B)* SLC4A7 expression (n=3), APP FL and CTF (n = 4), and LAMP1 (n = 4). Surface levels are normalised to control N-Cadherin whilst total levels are normalised to β-actin. Data expressed as a percentage of the relevant control shRNA. SNX27 and VPS35 shRNA data analysed by an one-way ANOVA followed by a post hoc Dunnett’s test compared to Control 1. SNX3+SNX12 shRNA data analysed by an unpaired t-test compared to Control 2. Error bars represent mean ± SEM. ****, p ≤ 0.0001; ***, p ≤ 0.001; **, p ≤ 0.01; ***, *, p ≤ 0.05; ns, not significant.

**Supplementary Figure 3.**
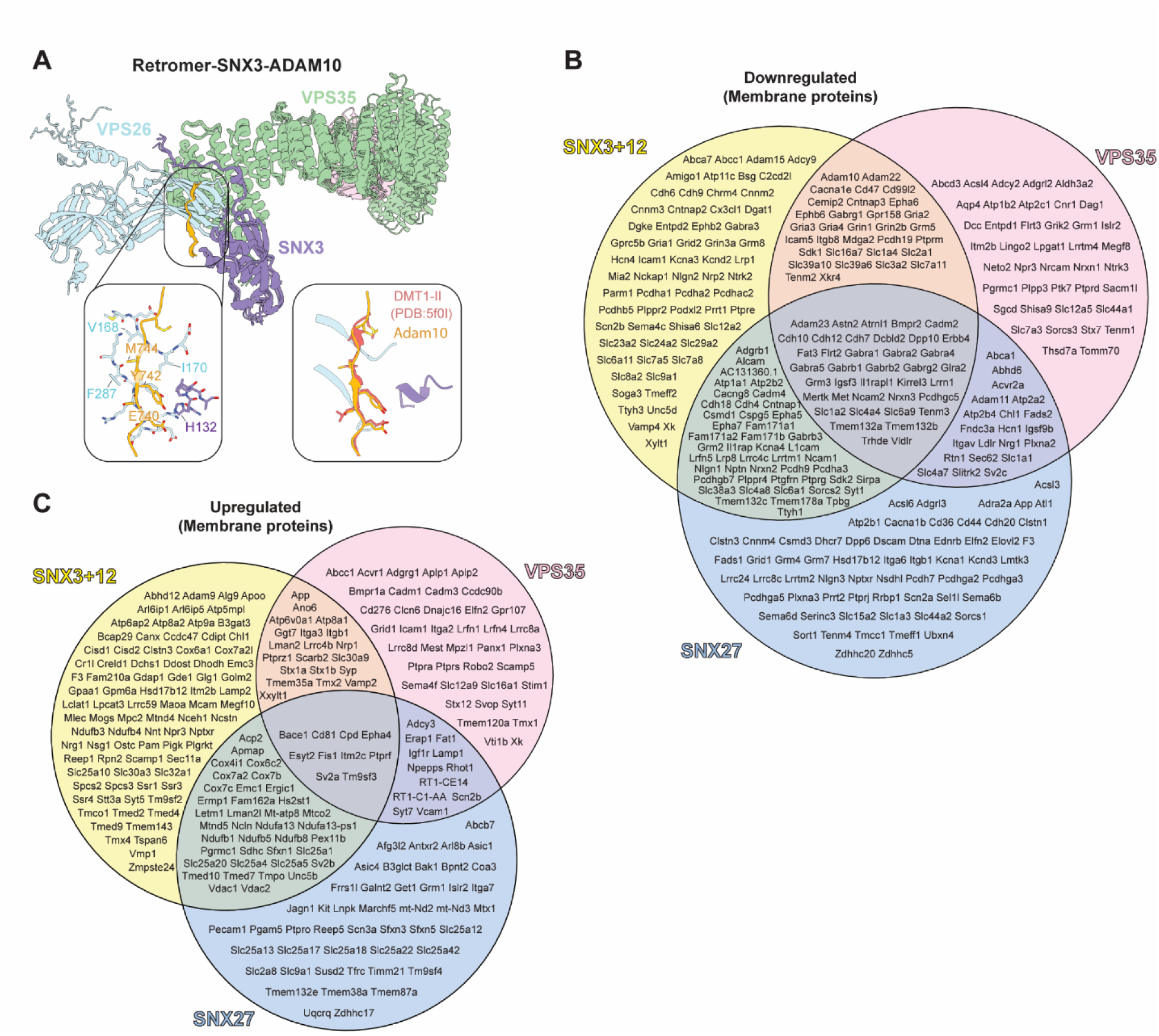
**(A)** AlphaFold3 predicted model of SNX3-Retromer binding to the cytoplasmic tail of ADAM10, compared to the published structure bound to DMT1-II (PDB 5f0I). **(B-C)** Venn diagrams displaying membrane proteins with significantly (**B**) downregulated (log_2_ fold change < −0.263) or (**C**) upregulated (log_2_ fold change > 0.263) surface abundance across SNX3+SNX12 (yellow), VPS35 (pink) and SNX27 (blue) depleted rat cortical neurons (p < 0.05), relative to control non-targeting shRNAs. Primary gene names as listed on Uniprot are shown.

**Supplementary Figure 4.**
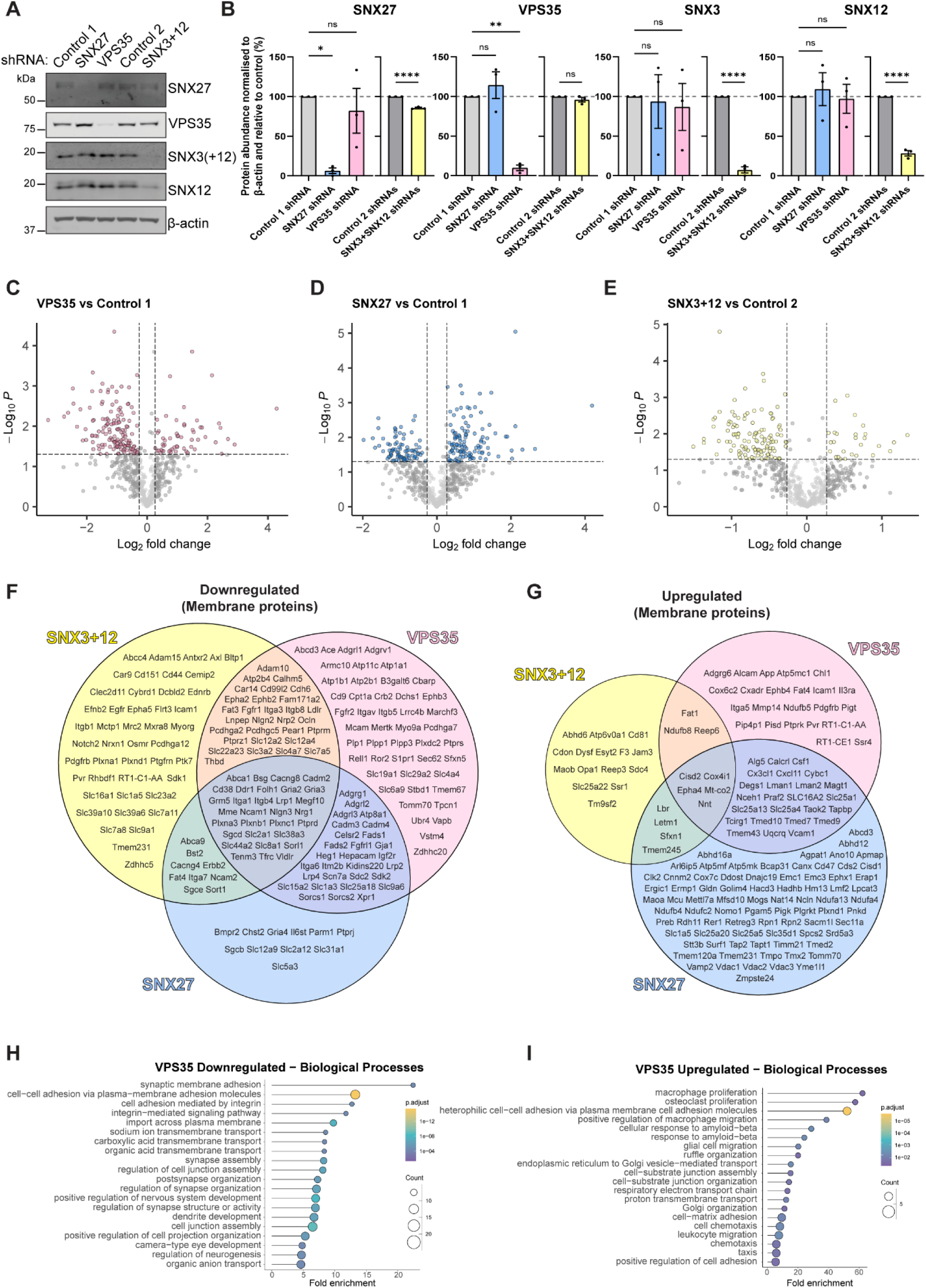
(**A**) Fluorescence-based western blot of rat cortical astrocytes transduced with either non-targeting shRNAs (Control 1 and Control 2) or shRNAs targeting SNX27, VPS35 or SNX3+SNX12, blotted for endogenous SNX27, VPS35, SNX3, SNX12 and β-actin. (**B**) Quantification of immunoblots shown in *(A)* across 3 independent experiments (n = 3). Data expressed as a percentage of the relevant control shRNA. SNX27 and VPS35 shRNA data analysed by an one-way ANOVA followed by a post hoc Dunnett’s test compared to Control 1. SNX3+SNX12 shRNA data analysed by an unpaired t-test compared to Control 2. Error bars represent mean ± SEM. ****, p ≤ 0.0001; **, p ≤ 0.01; ***, *, p ≤ 0.05; ns, not significant. (**C**-**E**) Volcano plots of membrane proteins in TMT surface proteomes of (**C**) VPS35, (**D**) SNX27 and (**E**) SNX3/SNX12 suppressed rat cortical astrocytes compared to non-targeting shRNA controls (Control 1 or Control 2). Dashed lines depict significance thresholds p ≤ 0.05 and log_2_ fold change ± 0.263. (**F**-**G**) Venn diagrams displaying membrane proteins with significantly (**F**) downregulated (log_2_ fold change < −0.263) or (**G**) upregulated (log_2_ fold change > 0.263) surface abundance across SNX3/SNX12 (yellow), VPS35 (pink) and SNX27 (blue) depleted rat cortical astrocytes (p < 0.05) relative to control non-targeting shRNAs. Primary gene names as listed on Uniprot are shown. **(H)** Downregulated or **(I)** upregulated surface proteins in VPS35 suppressed rat cortical neurons converge into biological pathways as shown by gene ontology analyses using Cluster Profiler.

**Supplementary Figure 5.**
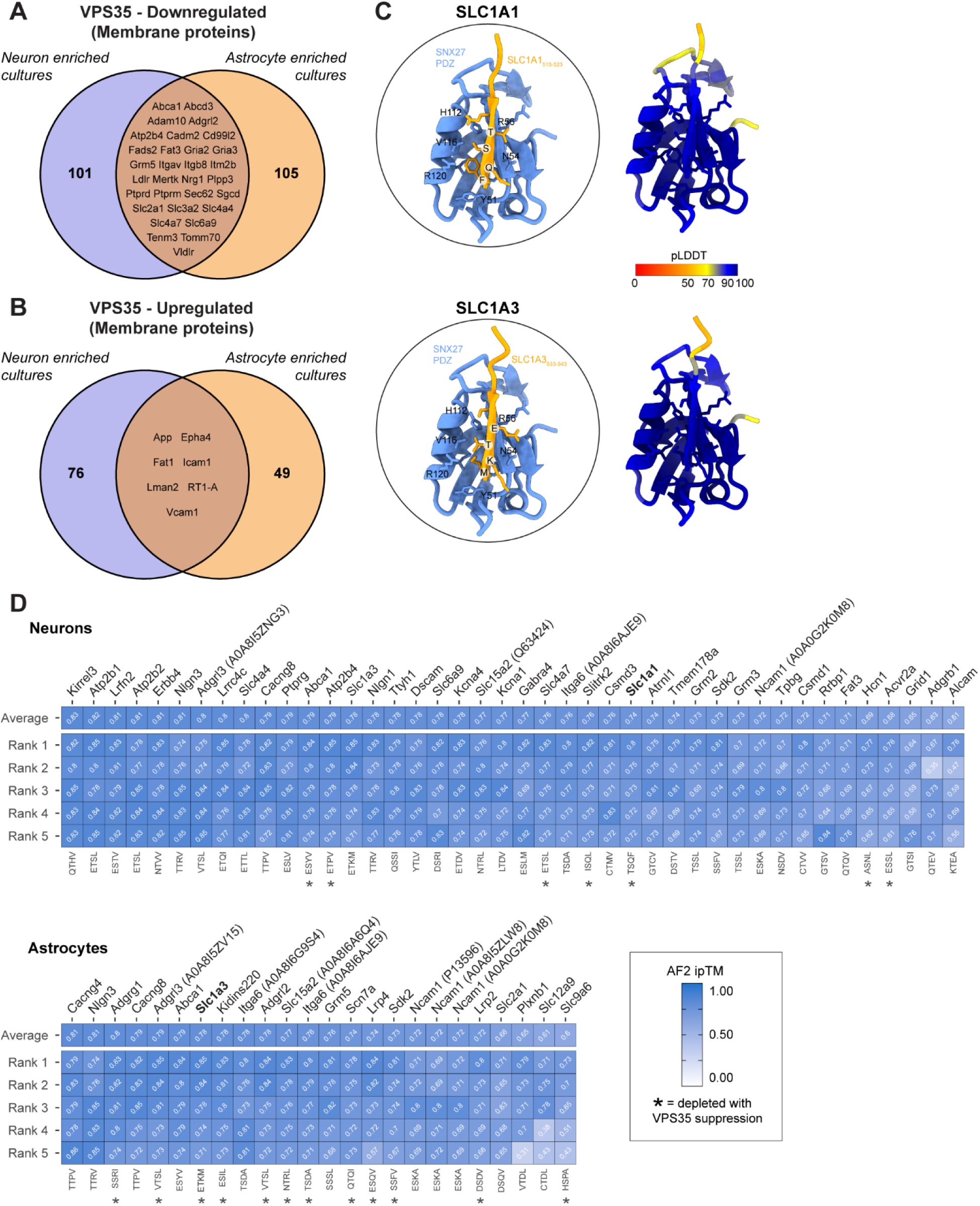
(**A**-**B**) Venn diagrams comparing membrane proteins significantly (**A**) downregulated (log_2_ fold change < - 0.263) and (**B**) upregulated (log_2_ fold change > 0.263) in rat cortical neuron (purple) and astrocyte (orange) enriched cultures with VPS35 suppression relative to non-targeting shRNA control (p < 0.05). For overlapping proteins, primary gene names as listed on Uniprot are shown. (**C**) AlphaFold2 predicted model of SNX27 interacting with the PDZbm in the cytoplasmic tail of SLC1A1 and SLC1A3. (**D**) AlphaFold2 identified 42 neuronal cargoes and 22 astrocytic cargoes that contain a PDZbm with an average interface predicted template modelling score (iPTM) higher than 0.6 against SNX27-PDZ predicting high confidence binding. The glutamate transporters SLC1A1 and SLC1A3 are highlighted in bold.

## Supplementary Table legends

**Supplementary Table 1**

Surface proteomics data from DIV19 rat cortical neurons transduced with either control non-targeting shRNAs (Control 1 and 2) or shRNAs against SNX27, VPS35 or SNX3/SNX12. Data is displayed for each protein raw and normalised to total peptide abundance, analysed as paired t-tests, with membrane and subcellular location information obtained from Uniprot. Log_2_ fold change and P-values from the paired t-test on normalised data, filtered for proteins labelled as membrane and non-contaminants, were used for further analysis as shown in pages 2-4.

**Supplementary Table 2**

Comparison of proteins with significantly altered surface abundance with VPS35 suppression compared to a non-targeting shRNA control (Control 1) in rat cortical neurons from the current manuscript, with significant hits from previously published surface proteomics datasets with VPS35 depletion in HeLa and H4 cells (Daly et al., 2023; Steinberg et al., 2013).

**Supplementary Table 3**

Comparison of proteins with significantly altered surface abundance with SNX27 suppression compared to a non-targeting shRNA control (Control 1) in rat cortical neurons from the current manuscript, with significant hits from previously published datasets: surface proteomics with SNX27 depletion in HeLa cells and SNX27 interactome in rat cortical neurons (McMillan et al., 2021; Steinberg et al., 2013).

**Supplementary Table 4**

AlphaFold2 modelling of neuronal SNX27-dependent cargoes that contain a PDZbm with the SNX27 PDZ domain. The final 10 amino acids at the carboxy-terminal of cytoplasmic tails were modelled. Data shows the mean interface predicted template modelling score (iPTM) across 5 ranks. Scores higher than 0.6 predict high confidence binding.

**Supplementary Table 5**

Confidence metrics from AlphaFold2 and AlphaFold3 modelling of SNX3-Retromer with the top 25 SNX3/SNX12 dependent single-pass transmembrane proteins in neuronal cultures. Data from AlphaFold3 modelling using full length cargo proteins and cytoplasmic tails alone shown as average iPTM score across 5 ranks, as well as the sum of these averages across both. pDockQ2 and interfacial iPTM scores of AlphaFold2 analysis with cytoplasmic tails also shown.

**Supplementary Table 6**

Surface proteomics data from rat cortical astrocytes transduced with either control non-targeting shRNAs (Control 1 and 2) or shRNAs against SNX27, VPS35 or SNX3/SNX12. Data is displayed for each protein raw and normalised to total peptide abundance, analysed as paired t-tests, with membrane and subcellular location information obtained from Uniprot. Log_2_ fold change and P-values from the paired t-test on normalised data, filtered for proteins labelled as membrane and non-contaminants, were used for further analysis as shown in pages 2-4.

**Supplementary Table 7**

AlphaFold2 modelling of astrocytic SNX27-dependent cargoes that contain a PDZbm with the SNX27 PDZ domain. The final 10 amino acids at the carboxy-terminal of cytoplasmic tails were modelled. Data shows the mean interface predicted template modelling score (iPTM) across 5 ranks. Scores higher than 0.6 predict high confidence binding.

**Supplementary Table 8**

Confidence metrics from AlphaFold2 and AlphaFold3 modelling of SNX3-Retromer with the top 25 SNX3/SNX12 dependent single-pass transmembrane proteins in astrocyte cultures. Data from AlphaFold3 modelling using full length cargo proteins and cytoplasmic tails alone shown as average iPTM score across 5 ranks, as well as the sum of these averages across both. pDockQ2 and interfacial iPTM scores of AlphaFold2 analysis with cytoplasmic tails also shown.

## Notes

### Competing Interest Statement

The authors have declared no competing interest.

